# Comparative genomics of tarakihi (*Nemadactylus macropterus*) and five New Zealand fish species: assembly contiguity affects the identification of genic features but not transposable elements

**DOI:** 10.1101/2022.08.01.502366

**Authors:** Yvan Papa, Maren Wellenreuther, Mark A. Morrison, Peter A. Ritchie

**Author notes:** Corresponding author. Address: School of Biological Sciences, Victoria University of Wellington, PO Box 600, Wellington 6140, New Zealand).

## Abstract

Comparative analysis of whole-genome sequences can provide valuable insights into the evolutionary patterns of diversification and adaptation of species, including the genome contents and the regions under selection. However, such studies are lacking for fishes in New Zealand. To supplement the recently sequenced genome of tarakihi (*Nemadactylus macropterus*), the genomes of five additional percomorph species native to New Zealand (king tarakihi (*Nemadactylus* n.sp.), blue moki (*Latridopsis ciliaris*), butterfish (*Odax pullus*), barracouta (*Thyrsites atun*), and kahawai (*Arripis trutta)*) were determined and assembled using Illumina sequencing. While the proportion of repeat elements was highly correlated with the genome size (*R*^2^ = 0.97, *P* < 0.01), most of the metrics for the genic features (e.g. number of exons or intron length) were significantly correlated with assembly contiguity (| *R*^2^| = 0.79–0.97). A phylogenomic tree including eight additional high-quality fish genomes was reconstructed from sequences of shared gene families. The radiation of Percomorpha was estimated to have occurred c. 112 mya (mid-Cretaceous), while the Latridae have diverged from true Perciformes c. 83 mya (late Cretaceous). Evidence of positive selection was found in 65 genes in tarakihi and 209 genes in Latridae: the largest portion of these are involved in the ATP binding pathway and the integral structure of membranes. These results and the *de novo* genome sequences can be used to (1) inform future studies on both the strength and shortcomings of scaffold-level assemblies for comparative genomics and (2) provide insights into the evolutionary patterns and processes of genome evolution in bony fishes.

## Introduction

The analyses of DNA sequences have enabled an extensive number of studies to be conducted into the evolutionary history of organisms. These have been used to investigate the underlying evolutionary mechanisms at both the inter-population and inter-species levels. While DNA studies have previously only been carried out on small regions of the genome (e.g. short mitochondrial or nuclear DNA sequences via Sanger sequencing), recent advances in sequencing technologies have greatly improved the acquisition of large, genome-wide DNA sequence data sets. This technical advance enabled the field of comparative genomics to rapidly expand (Hardison, 2003; Miller et al., 2004). At its core, comparative genomics utilizes a range of analyses that align contiguous sequences of long stretches of the genome to identify orthologous regions (i.e. sequences that share a common ancestor) and quantify the amount and type of change that has occurred between them (Ellegren, 2008). Genomic sequence similarities and dissimilarities allow inferences to be made about gene functions and structural variation, and how these might influence the evolutionary process. One of the key goals in the field of molecular evolution is to elucidate how selection operates in the genome (Nielsen, 2005). By minimizing the stochastic effects seen in short sequences and variation among different genes, genomics make it possible to detect signatures of selection in specific regions of the genome (Ellegren, 2008; Vitti et al., 2013). This is typically achieved by testing for deviations from neutral expectations (Zhang, 2005).

Genome assembly and annotation allow for the discovery, description, and comparison of orthologous gene coding regions and other genomic features that play key roles in the evolutionary history of organisms. For example, it is possible to detect and characterize repeat elements (RE) in the genome (Lerat, 2018). REs can either consist of tandem repeats (e.g. satellite DNA, also referred to as “simple” repeats) or interspersed repeats (Lerat, 2018; Richard et al., 2008). Interspersed repeats include tRNA genes, genes paralogues, and transposable elements (TE) (Richard et al., 2008). TEs are stretches of DNA that have the capacity to move from one position to another along the chromosomes. There are several types of TEs, classified according to their transposition intermediate (RNA or DNA), their structural features, and their evolutionary origin (Wicker et al., 2007). There is a growing number of evidence that REs make a significant contribution to genome evolution (Biémont, 2010; Biémont & Vieira, 2006) and can have both deleterious and beneficial effects (Chuong et al., 2017). TEs can drive genetic diversification by e.g. altering the protein coding capacity of genes, inducing structural rearrangement, and providing new material on which natural selection can act on. Moreover, both TEs and simple repeats can have an influence on the size of genomes (Lerat, 2018; Sotero-Caio et al., 2017; Z. Yuan et al., 2018). However, very few studies have used genome-wide data to compare the proportion and diversity of repeat elements in several teleost fishes genomes (Brawand et al., 2014; Gao et al., 2016; Shao et al., 2019).

There are other genomic features that are seldom explored. This includes “genic features”, such as the number and diversity metrics of genes and their components (e.g. exons, introns, UTR regions). There is still much to discover about these genic features from a comparative genomics perspective. For example, a synthetic review of the genome content of eukaryotes found that while the number of genes increases with genome size, the proportion of coding elements decreases and the proportion of introns increases (Elliott & Gregory, 2015). It was also suggested that there might be a positive relationship between the exonization of TEs and intron length in animals (Sela et al., 2010). However, few genome assembly studies report these genic features metrics and usually report only the number of genes. Elliott & Gregory (2015) emphasized that the lack of reporting of metrics like the number and proportion of coding regions and introns is an issue that should be addressed. A recent study that explored the patterns of size, GC content, number of chromosomes and number of genes in fish genomes (Randhawa & Pawar, 2021) highlighted that “surprisingly, no study exists on record that has used the WGS annotation data to defines the trends, effects of taxonomic distribution and interrelation of genome attributes”. Yet, such studies have successfully unveiled inter-lineage genomic characteristic patterns in e.g. rodents (Capilla et al., 2016) and birds (Feng et al., 2020).

Current sequencing technologies (e.g. long reads of thousands of bp and scaffolding data like Hi-C) allow for chromosome level and phased assemblies to anchor the totality of genomic information (e.g. number of chromosomes, structural variation) and minimize the risks of incorrect inferences (e.g. unmerged haplotigs, potential scaffolding errors). However, the required integrity of DNA can be hard to obtain from sampling wild-caught specimens, particularly when the sampling conditions prevent the rapid conservation of the tissue samples in optimal conditions. Rapid degradation of DNA occurs in the first few hours after harvesting a specimen (Oosting et al., 2020). This is particularly problematic for producing long-read sequences (Klingström et al., 2018) and keeping the chromatin integrity for Hi-C data. Depending on the application, genomes of sufficient quality can be obtained with less sampling constraints using only short-read (100–1000 bp) technology, for which partial degradation of DNA is less of an issue. While genome assemblies based on short reads only will invariably be fragmented to hundreds or thousands of scaffolds, sufficient read coverage can still lead to high-quality contigs and scaffolds that can be used to carry out diverse comparative genomic analyses (Feng et al., 2020; Malmstrøm et al., 2016, 2017).

Although more than a thousand species of fishes occur in New Zealand waters (Roberts et al., 2020) and its marine ecosystem is highly valuable commercially, recreationally and culturally, genome-wide analyses have seldom been used to study its ichthyofauna (Papa, Oosting, et al., 2021). The few exceptions have focused on genomic variation at the intra-species and population level (Catanach et al., 2019; Koot et al., 2021; Oosting, 2021). The main goal of this study was to investigate the patterns of genome-wide variation and evolution of tarakihi (*Nemadactylus macropterus*) and five newly sequenced New Zealand fish species (king tarakihi (*Nemadactylus* n.sp.), blue moki (*Latridopsis ciliaris*), greenbone butterfish (*Odax pullus*), barracouta (*Thyrsites atun*), and kahawai (*Arripis trutta*) (Figure 1) in a comparative genomic framework. To achieve this, the five new genomes were assembled using Illumina short read sequences. The high-coverage genome assembly and annotation of tarakihi produced in Papa, Wellenreuther, et al. (2021) was combined with these five genomes to explore the diversity of repeat elements and genic features. High-quality genomic data from eight other fishes retrieved from Ensembl were then added to identify genes under positive selection in tarakihi and Latridae in a phylogenomic framework.

**Figure 1.**
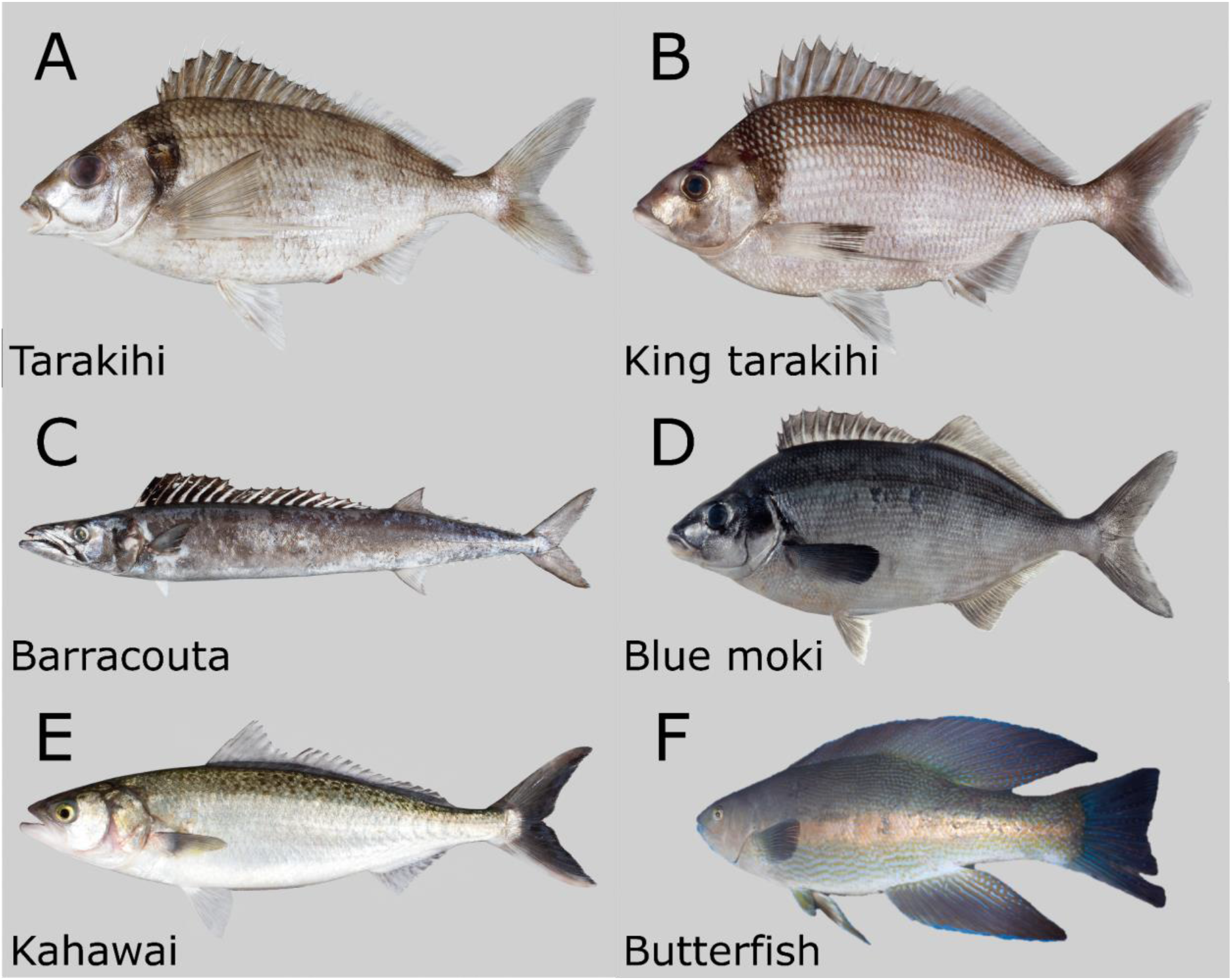
The six fish species for which a *de novo* genome assembly has been produced in Papa, Wellenreuther, et al. (2021) (A) and this study (B–F). (A) Tarakihi (*Nemadactylus macropterus*: Centrarchiformes, Latridae) (B) King tarakihi (*Nemadactylus* n.sp.: Centrarchiformes, Latridae) (C) Barracouta (*Thyrsites atun*: Scombriformes, Gempylidae) (D) Blue moki (*Latridopsis ciliaris*: Centrarchiformes, Latridae) (E) Kahawai (*Arripis trutta*: Scombriformes, Arripidae) (F) Greenbone butterfish (*Odax pullus*: Labriformes, Labridae). Pictures from Roberts et al. (2015), courtesy of the Museum of New Zealand Te Papa Tongarewa.

## Materials and Methods

### Tissues collection and DNA extraction

DNA was isolated from five specimens each belonging to a different fish species (king tarakihi, blue moki, butterfish, barracouta, and kahawai) for the respective *de novo* genome assemblies (Figure 1). The fish were sampled opportunistically, based on availability, as part of a sampling campaign for the genome assembly and population genomics of tarakihi (Papa, Morrison, et al., 2021; Papa, Wellenreuther, et al., 2021). The king tarakihi specimen was collected by a commercial fishing trawler around Three King Islands at a depth between 140 and 250 m. The standard length was 477 mm, weight was 1,900 g, and it was identified as a female by observation of the gonads. A piece of muscle tissue (c. 2 cm) from the tail was collected and stored in 99% EtOH at −20 °C. The barracouta (70 cm, male) was captured by a recreational fisherman in the Wellington harbour (New Zealand). The three remaining specimens (blue moki, butterfish, and kahawai, sex undetermined) were caught by recreational spear-fishers off Island Bay, around the south coast of Wellington. For these and the barracouta, a piece of the pectoral fin was collected and immersed in 20% DMSO, 0.25 M EDTA, NaCl saturated solution (DESS) and stored at −20 °C. Total genomic DNA was extracted from all tissues with a modified high-salt extraction protocol (Aljanabi & Martinez, 1997) that included an RNase step. The extracted DNA was re-suspended in Tris-EDTA buffer (10 mM Tris-HCl pH 8.0, 0.1 mM EDTA). The purity of DNA samples was assessed using a NanoPhotometer^®^ NP80 (Implen). The high-molecular-weight DNA was visualised using ethidium bromide-straining gel electrophoresis and the quantity of double-strand DNA was measured using a Qubit™ dsDNA BR Assay Kit.

### Library preparation and sequencing

Prior to sequencing, the genome sizes of the five fish species were estimated based on previously published studies to be approximately 800 Mb (+-c. 200 Mb), which corresponds to the average genome size of bony fishes (Fan et al., 2020) and the approximate genome size for most percomorphs (Z. Yuan et al., 2018). This estimation was rounded up to a less conservative 1 Gb and was used to determine the required sequencing coverage. Coverage depth needed for short Illumina reads was estimated to be 20–30×, although a plateau in BUSCO completeness is often reached at around 15× (Malmstrøm et al., 2017). Samples were whole-genome sequenced with the aim of obtaining at least 25 Gb per sample.

The purified DNA samples were sent to the Australian Genome Research Facility (AGRF, Melbourne, Australia) for DNA library preparation and sequencing. Illumina DNA shotgun libraries were prepared following the Nextera DNA FLEX low volume protocol with Nextera DNA Combinatorial Dual Indexes (Illumina) for fragment sizes 300–350 bp. Short reads sequencing of 150 bp paired-end reads was performed on NovaSeq 6000 (Illumina) with NovaSeq 6000 S4 Reagent Kit and NovaSeq XP 4-Lane Kit for 300 cycles. Sequencing was performed during several rounds on different lanes depending on the well availability during sequencing of other species (tarakihi and snapper (*Chrysophrys auratus)*) from other studies. Base calling and quality scoring were performed with RTA3 software v3.3.3. De-multiplexing of the sequencing data was performed using Illumina bcl2fastq pipeline v2.20.0.422.

### Quality and contamination filtering

Quality of reads was assessed and visualised at several steps on the filtering pipeline using FastQC v0.11.7 (Andrews, 2018) and results were compiled in MultiQC v1.7 (Ewels et al., 2016). Raw paired-end reads were filtered as follows: First, reads that contained adapter contamination were discarded. For this, Trimmomatic v0.39 (Bolger et al., 2014) was used with parameter `NexteraPE-PE.fa:2:30:10` to trim adapters contamination from the reads. seqkit v0.13.2 (Shen et al., 2016) was used to keep only the reads that were not trimmed (i.e. reads with a length of 150bp). Then, reads that contained more than 10% uncertain bases (i.e. Ns) and/or for which the proportion of bases with Quality Value ≤ 19 was over 50% were also filtered out using custom bash scripts. Paired-end reads were both discarded if either the forward or the reverse read did not pass the above filtering criteria. Finally, DNA sequence contamination from archaea, bacteria, viruses or human DNA was detected and filtered out using Kraken v2.0.7-beta (Wood et al., 2019) with the MiniKraken2 v2 8GB database (Wood, 2019).

### Mitogenome assembly and exclusion

For each fish sample, quality-filtered and contaminant-free Illumina reads were mapped against the complete mitochondrial sequence of an available close species with Geneious v11.04 (Kearse et al., 2012) by running five iterations of the default mapper set to highest sensitivity. The mitochondrial genomes used as references and their Genbank accession numbers were the following. Barracouta: silver gemfish, *Rexea solandri* (NC_023952.1); blue moki: Peruvian morwong, *Cheilodactylus variegatus* (KP704218.1), butterfish: herring cale, *Olisthops cyanomelas* (NC_009061.1), kahawai: same species, *Arripis trutta* (NC_015787.1). For the king tarakihi, the tarakihi (*Nemadactylus macropterus*) mitogenome produced in Papa, Wellenreuther, et al. (2021) was used as reference. The assembled mitogenomes were then annotated using the MitoAnnotator web interface (Iwasaki et al., 2013). Sequences that were of mitochondrial origin were then filtered out as follows: bwa-kit v0.7.15 (Li & Durbin, 2009) was first used to align the Illumina reads to their respective indexed reference mitogenomes with default parameters. All the reads from the alignment that did not map to the mitogenome were then extracted into a new mitochondria-free alignment using SAMtools v1.9 (Li et al., 2009) `view` with parameters `-b -f 4`, sorted by name and finally converted back to FASTQ paired-end reads with bedtools v2.27.1 (Quinlan & Hall, 2010).

Once the mitogenomes were assembled, some specific sections were selected for BLAST analysis on NCBI to rule out the possibility of incorrect morphology-based identification of the specimens at collection or DNA sample contamination with another fish species. A c. 550 bp subset of the cytochrome *c* oxidase I region of barracouta, blue moki, and kahawai were blasted against the NCBI nucleotide database. In all cases, the best matches were the correct species with percent identity scores between 99.82% and 100%, consistent with the morphological identification of the species used in each sample. Since there was no COI sequence available for the butterfish on NCBI, the 16S RNA region was used instead and showed a 100% identity match. The king tarakihi specimen was not blasted because there were no sequences available for this species on NCBI. The specimen was already identified with high confidence using morphology and based on the phylogenetic position of the sequence data (see Papa, Morrison, et al. (2021)).

### Genome assembly

Genome assemblies were performed with the Maryland Super-Read Celera Assembler, MaSuRCA v3.4.1 (Zimin et al., 2013, 2017). All assemblies were run with recommended parameters for medium-size genomes based on short paired-end Illumina reads only (PE = pe 350 50, EXTEND_JUMP_READS = 0, GRAPH_KMER_SIZE = auto, USE_LINKING_MATES = 1, USE_GRID = 0, GRID_BATCH_SIZE = 300000000, LHE_COVERAGE=25, MEGA_READS_ONE_PASS=0, LIMIT_JUMP_COVERAGE = 300, CA_PARAMETERS = cgwErrorRate = 0.15, CLOSE_GAPS = 1, NUM_THREADS = 32, SOAP_ASSEMBLY = 0). The `JF_SIZE` parameter was set as 20^9^ for king tarakihi, 13^9^ for barracouta and blue moki, and 10^9^ for butterfish and kahawai.

Assemblies scaffolds were sorted by size using seqkit v0.13.2 and renamed with simple numbers (i.e. “1” for the longest scaffold, then “2”, etc.) with command `replace –p .+ –r “{nr}”`. Basic contiguity statistics of all assemblies were computed with bbmap v38.31 (Bushnell, 2018) script `stats.sh`. Genome completeness of each assembly was assessed with the Benchmarking Universal Single-Copy Orthologs (BUSCO) tool v3.0.2 (Simão et al., 2015) and its dependencies (Augustus v3.3.1 (Stanke et al., 2004), NCBI blast+ v2.7.1 (Camacho et al., 2009), hmmer v3.2.1 (Eddy, 2011), and R v3.6.0 (R Core Team, 2020)). Contiguity and completeness of the genome assemblies were graphically visualised with assembly-stats v17.02 (Challis, 2017) as implemented in the grpiccoli container (Piccoli, 2021).

### Genome annotation

Repetitive elements in the five genomes were identified using the same method and tools as reported in Papa, Wellenreuther, et al. (2021). During the repeat identification step, the Actinopterygii homology-based repeat library produced in Papa, Wellenreuther, et al. (2021) was combined with each of the *de novo* repeat libraries produced separately for each species. Genome annotation was carried out with the MAKER v2.31.10 (Holt & Yandell, 2011) pipeline on the unmasked genomes. Before annotation, the simple repeats were filtered out of the repeats annotation file. This allowed to hard-mask only the complex repeats regions and keep the simple repeats available for soft-masking by MAKER (see method details in Papa, Wellenreuther, et al. (2021)). For each assembly, the first round of MAKER was run on the unmasked genome using protein and transcriptome data for gene models prediction (`protein2genome=1, est2genome=1`) and the GFF of complex repeats only for hard masking (model_org=`simple`). Protein sequences of green spotted puffer (*Tetraodon nigroviridis*), Japanese puffer (*Takifugu rubripes*), medaka (*Oryzias latipes*), Nile tilapia (*Oreochromis niloticus*), southern platyfish (*Xiphophorus maculatus*), spotted gar (*Lepisosteus oculatus*), three-spined stickleback (*Gasterosteus aculeatus*), and zebrafish (*Danio rerio*) were downloaded from Ensembl release version 103 (Kersey et al., 2016) and used as protein homology in the first round. The transcriptome data used differed depending on the species but was always sourced from a reasonably closely related species (parameter `altest` was set instead of `est`): for the king tarakihi and the blue moki, the repeat-filtered, non-redundant Iso-Seq transcripts produced in Papa, Wellenreuther, et al. (2021) from the tarakihi were used. The Transcriptome Shotgun Assembly (TSA) dataset from the Atlantic mackerel *Scomber scombrus* (Prefix ID: GHRT01, length: 379.6 Mb) was downloaded from NCBI and used for the barracouta and the kahawai. Similarly, the TSA dataset from the hogfish *Lachnolaimus maximus* (Prefix ID: GFXS01, length: 304.2Mb) was used for the butterfish. Training files for the *ab initio* gene predictors Augustus v3.3.1 (Stanke et al., 2004) and SNAP v2013.11.29 (Korf, 2004) were generated based on round 1 results. The second round of MAKER, the gene model set quality control step, and the functional annotation were carried out exactly like in Papa, Wellenreuther, et al. (2021), except that the GFF protein alignment produced in round 1 was used as evidence (`protein2genome=0`) for round 2. The influence of both genome size and genome assembly fragmentation on the annotation of several genomic features was tested using Pearson correlation tests with R v4.02 (R Core Team, 2020) base function `cor.test`. The genomic features included e.g. the proportion of repeat elements, the number of genes or the total intron length. The BUSCO completeness score was used to represent genome contiguity as it is a strong predictor of this metric (Jauhal & Newcomb, 2021).

### Gene family identification and phylogenetic tree construction

Protein and DNA coding sequences of each of the six New Zealand species were retrieved from their respective MAKER annotation pipeline. Protein and DNA coding sequences of zebrafish (*Danio rerio*), three-spined stickleback (*Gasterosteus aculeatus*), spotted gar (*Lepisosteus oculatus*), Nile tilapia (*Oreochromis niloticus*), medaka (*Oryzias latipes*), Japanese puffer (*Takifugu rubripes*), green spotted puffer (*Tetraodon nigroviridis*), and southern platyfish (*Xiphophorus maculatus*) (i.e. the same species that were used for the homology-based genome annotation) were downloaded from Ensembl release v103 (Kersey et al., 2016). Single-copy gene family orthologs between the 14 species were found using OrthoFinder v2.5.2 (Emms & Kelly, 2019) on the protein sequences with default parameters. The protein sequences in each orthogroup were then aligned with MAFFT v7.480 (Katoh & Standley, 2013). Poorly aligned positions and divergent regions were removed with Gblocks v0.91b (Castresana, 2000). All alignments were concatenated in one single alignment using seqkit v0.13.2 and the best substitution model was inferred with ModelTest-NG v0.1.7 (Darriba et al., 2020). The phylogenomic tree of the 14 species was built with MrBayes v3.2.7 (Ronquist et al., 2012) using the JTT+I+G4+F model with 20,000 generations and the default parameters (three heated chains, one cold chain and a burn-in of 25%). After calibrating some of the nodes with the interval values retrieved from TimeTree (www.timetree.org), the divergence times along the tree were calculated with MCMCTree from PAML v4.9 (Yang, 2007; Yang & Rannala, 2006) using the method for approximate likelihood with protein data. The gene family expansions and contractions along the tree branches were calculated with CAFE v4.2.1 (De Bie et al., 2006). The gene families used for the CAFE analysis were based on the orthogroups defined earlier by OrthoFinder, except that orthogroups were not used if they contained species with more than 100 gene copies due to issues with high variance.

### Tests for selection

Genes were investigated for evidence of positive selection separately in two lineages: the tarakihi and the Latridae (tarakihi, king tarakihi, and blue moki). Latridae were of particular interest because it is a speciose but taxonomically contentious clade (Kimura et al., 2018; Ludt et al., 2019) restricted to the Southern Hemisphere and no other genome assembly currently exists for this family. Selection was detected in the one-to-one orthologues as follows: the protein sequence alignments were converted to nucleotides codon alignments with PAL2NAL v14.1 (Suyama et al., 2006) using the corresponding DNA coding sequences. The alignments were polished with Gblocks v0.91b (parameter codon) and converted to phylip format with seqmagick v0.8.0 (https://github.com/fhcrc/seqmagick). Positively selected genes were detected with CodeML from PAML v4.9 with the branch-site model test, using one of the two selected lineages (tarakihi only or tarakihi, king tarakihi and blue moki) as the foreground branches. The null model assumed that the substitution rates at nonsynonymous and synonymous sites (d*N*/d*S* ratio) for all codons in all branches must be ≤1. The alternative model assumed that the foreground branches included codons evolving at d*N*/d*S* > 1, indicating the fixation of advantageous mutations. The analysis was applied on each gene (i.e. single-copy orthogroup) DNA alignment separately, using the tree topology obtained earlier with MrBayes. For each gene, the log likelihoods of the alternative (nL1) and null models (lnL0) were compared with a likelihood ratio test (ΔLRT = 2(lnL1 - lnL0)). The associated p-values were calculated under the chi-square distribution with 1 degree of freedom (df = 1). All p-values were then adjusted for multiple testing with the false discovery rate method. Genes were considered positively selected if the adjusted p-value was ≤ 0.05 and if at least one amino-acid site had a Bayes probability ≥ 95% of being positively selected. All Gene Ontology (GO) terms associated with the selected genes were then obtained from the UniProt database (Bateman, 2019). The zebrafish gene accession numbers were used for associated GO terms because this is a well-curated model species.

### General bioinformatics tools

Analyses were performed on the Victoria University of Wellington high-performance computer cluster Rāpoi. R analyses were performed in R v4.02 (R Core Team, 2020) on RStudio (RStudio Team, 2020). See the method section in Papa, Wellenreuther, et al. (2021) for more details on general bioinformatics tools and commands used.

## Results

### Genome sequencing and assembly

The DNA sequencing data sets contained 156–197 million reads after they were filtered for quality, contamination and mitochondrial sequences. This total amount corresponded to 23.43–29.52 Gb per species (Supplementary Table 1). The sizes of the five final genome assemblies (Figure 2, Table 1) ranged from 532 Mb (butterfish) to 714 Mb (barracouta). While the genome sizes of bony fishes cover a very wide range of values (0.34 Gb for *Tetraodon nigroviridis* to 2.97 Gb for *Salmo salar*), these results are concordant with the expected sizes for many percomorphs (Z. Yuan et al., 2018). Contiguity and completeness varied among species, with the number of scaffolds ranging from 58,102 (king tarakihi) to 150,595 (barracouta) and the N50s from 10,031 (blue moki) to 30,492 (king tarakihi). Based on the genome assembly sizes, the lowest read coverage was 33× (barracouta) and the highest was 52× (butterfish). King tarakihi, blue moki and kahawai coverages were all above 40×. Final genome completeness was also variable, with a BUSCO completeness for single-copy Actinopterygii orthologs ranging from 70.20% (kahawai) to 89.10% (king tarakihi). Both contiguity and completeness of the produced genomes were on par with quality metrics for fish assemblies using short Illumina reads (Malmstrøm et al., 2017).

**Figure 2.**
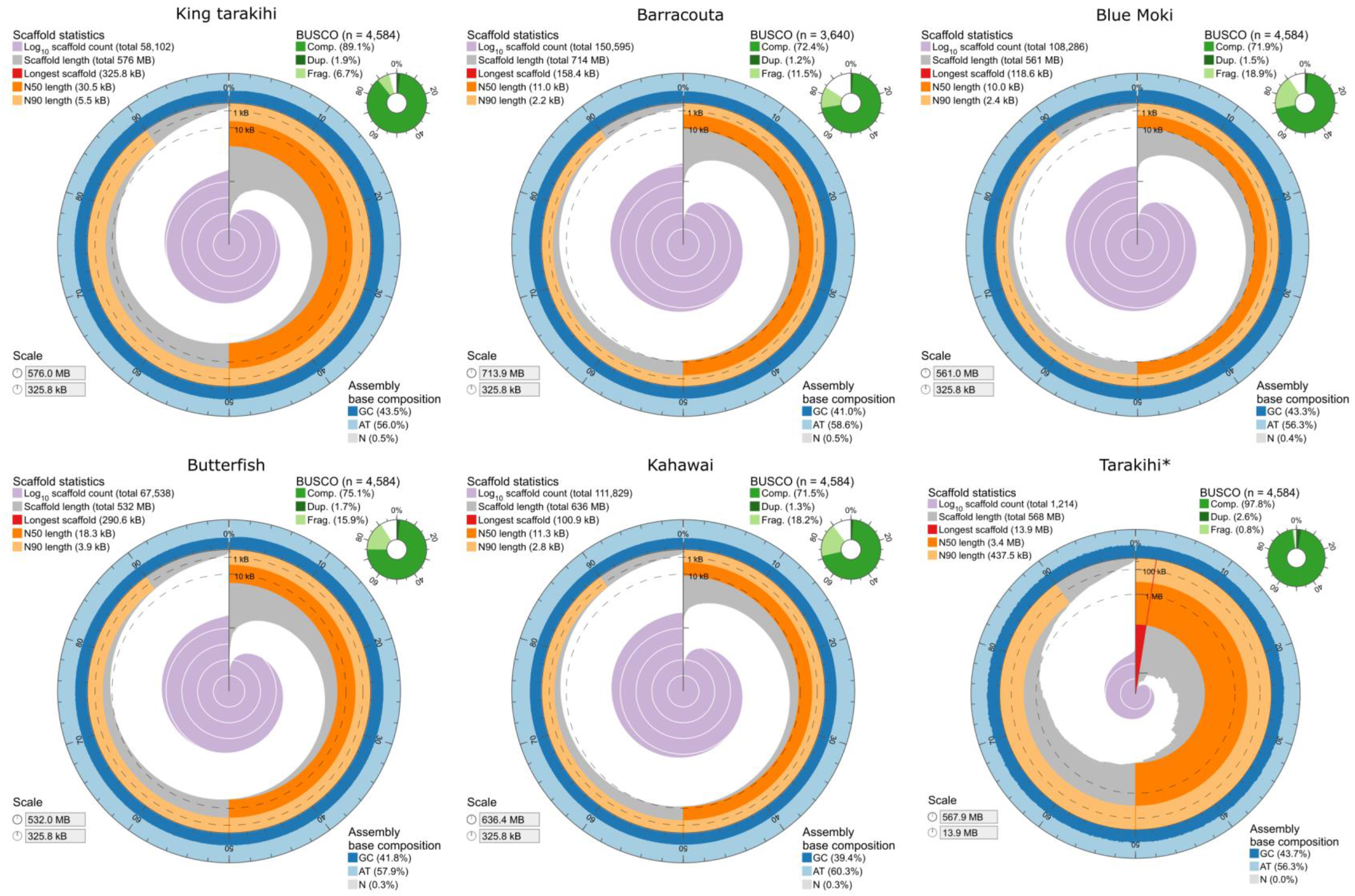
Visualisation of contiguity and completeness of the five new genome assemblies from this study and (*) the tarakihi assembly from Papa, Wellenreuther, et al. (2021). The contiguity is visualised in a circle representing the full assembly length (532–714 Mb). Lengths of longest scaffolds of the new assemblies ranged from 100.9 to 325.8 Kb. There were very few scaffolds (c. 2%) shorter than 100 Kb in length and the GC content was always uniform throughout.

**Table 1.** General statistics of the five new assemblies produced.

|  | King tarakihi | Barracouta | Blue moki | Butterfish | Kahawai |
| --- | --- | --- | --- | --- | --- |
| Genome Assembly |  |  |  |  |  |
| Scaffold assembly size (bp) | 575,828,338 | 713,861,692 | 560,782,505 | 531,871,215 | 636,431,267 |
| Reads coverage | 40.70× | 33.33× | 42.93× | 52.14× | 46.38× |
| Total number of scaffolds | 58,102 | 150,595 | 108,286 | 67,538 | 111,829 |
| Longest scaffold (bp) | 325,842 | 158,436 | 118,632 | 290,593 | 100,923 |
| Scaffold N50 (bp) / L50 | 30,492 / 5,169 | 10,964 / 17,677 | 10,031 / 15,899 | 18,273 / 7,941 | 11,325 / 16,202 |
| Proportion of gap sequences | 0.510% | 0.452% | 0.412% | 0.291% | 0.344% |
| Contigs size (bp) | 572,888,966 | 710,633,565 | 558,474,240 | 530,323,606 | 634,240,427 |
| Total number of contigs | 106,327 | 231,192 | 148,983 | 94,271 | 154,681 |
| Contig N50 (bp) / L50 | 12,009 bp / 13,187 | 5,899 / 33,702 | 6,943 / 23,019 | 11,807 / 12,336 | 7,607 / 24,285 |
| A / T / G / C bases (%) | 28.30 / 28.24 / 21.72 / 21.74 | 29.59% / 29.42% / 20.47% / 20.52% | 28.39% / 28.33% / 21.63% / 21.65% | 29.12% / 29.03% / 20.92% / 20.93% | 30.33% / 30.27% / 19.69% / 19.71% |
| Genome Completeness (4,584 Actinopterygii BUSCOs)<sup>†</sup> |  |  |  |  |  |
| Complete BUSCOs | 89.1% | 72.4% | 71.90% | 75.10% | 71.50% |
| Complete single-copy BUSCOs | 87.2% | 71.2% | 70.40% | 73.40% | 70.20% |
| Complete duplicated BUSCOs | 1.9% | 1.2% | 1.50% | 1.70% | 1.30% |
| Fragmented BUSCOs | 6.7% | 11.5% | 18.90% | 15.90% | 18.20% |
| Missing BUSCOs | 4.2% | 16.1% | 9.20% | 9.00% | 10.30% |
| Genome annotation |  |  |  |  |  |
| No. of protein-coding genes | 22,258 | 24,378 | 23,804 | 24,816 | 22,840 |
| % genes with AED < 0.5 | 92% | 86% | 87% | 85% | 96% |
| No. funct. annotated proteins | 21,650 | 23,840 | 23,269 | 24,266 | 22,570 |
| Mean gene length (bp) | 7,664 | 4,371 | 3,808 | 4,163 | 5,352 |
| Mean exon length (bp) | 160 | 170 | 173 | 169 | 166 |
| Mean intron length (bp) | 861 | 794 | 651 | 757 | 737 |

Note: See Papa, Wellenreuther, et al. (2021) for assembly statistics of the tarakihi. (†)Barracouta BUSCO completeness was assessed with Actinopterygii orthologs database v10 (3,640 BUSCOs) instead of v9 (4,584 BUSCOs) because of version software compatibility at the time of the analysis. The results from both orthologs databases are still comparable.

### Repetitive elements and genes

The total proportion of repetitive elements in the six genomes (Figure 3, Table 2, Supplementary Table 2) varied from 24.83% (butterfish) to 39.12% (barracouta). The proportion of repeat elements in the king tarakihi and the blue moki (c. 30%) is similar to the proportion obtained for the tarakihi (Figure 3, Table 2). These three species are from the same family (Latridae). The proportion of repeat elements was highly correlated to the genome size (Figure 3) (see Discussion).

**Figure 3.**
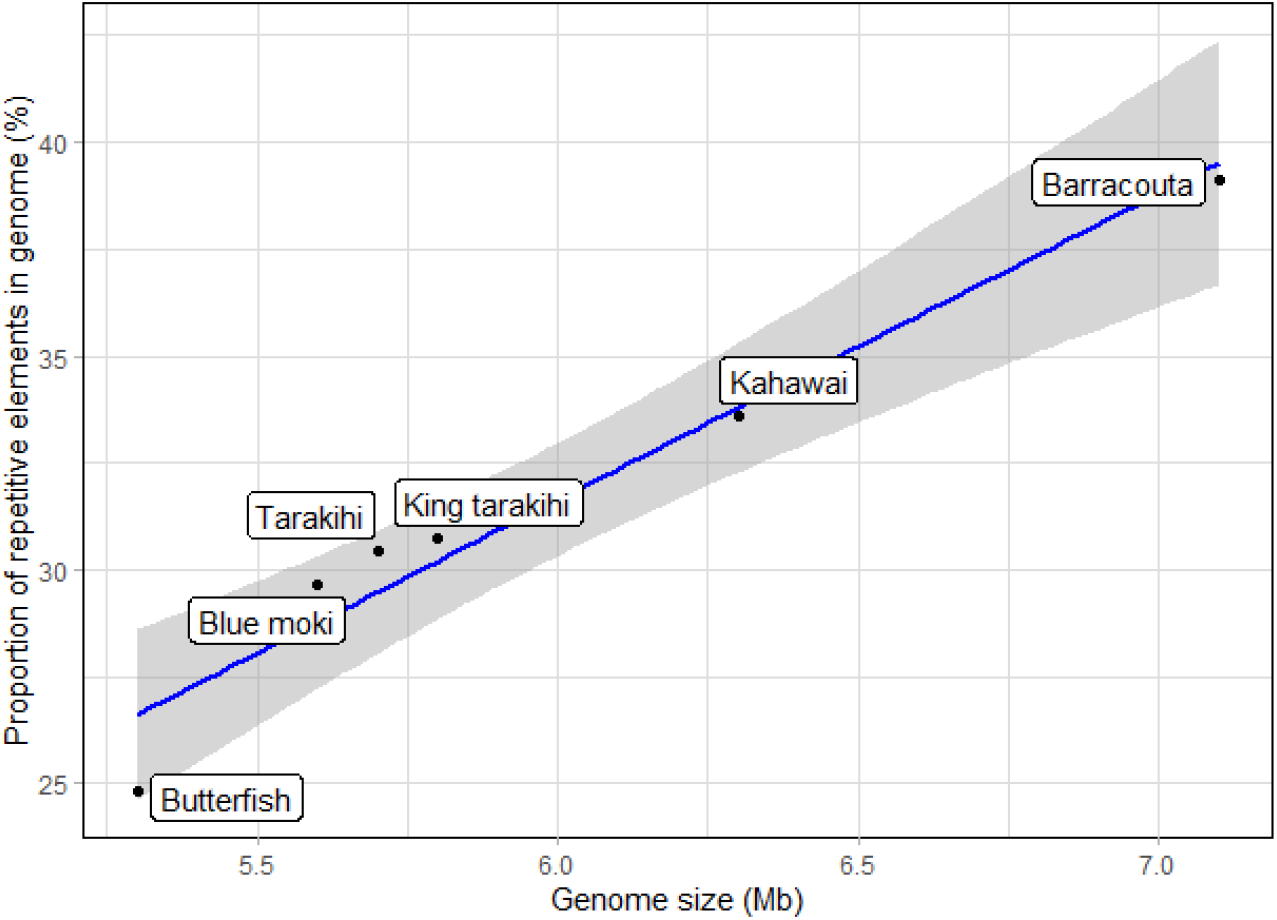
Correlation between genome sizes and the proportion of repetitive elements in the genome. Tarakihi, king tarakihi and blue moki are from the same family (Latridae).

**Table 2.** Gene features of the six fish genome assemblies

|  | Tarakihi | King tarakihi | Blue moki | Butterfish | Barracouta | Kahawai |
| --- | --- | --- | --- | --- | --- | --- |
| Genome length (bp) | 567,902,715 | 575,828,338 | 560,782,505 | 531,871,215 | 713,861,692 | 636,431,267 |
| Number of genes / mRNA / CDS | 20,169 | 22,258 | 23,804 | 24,816 | 24,378 | 22,840 |
| Number of exons | 217,298 | 185,789 | 129,362 | 131,762 | 130,618 | 154,018 |
| Number of introns | 197,129 | 163,531 | 105,558 | 106,946 | 106,240 | 131,178 |
| Mean exons per gene / mRNA | 11 | 8 | 5 | 5 | 5 | 7 |
| Total gene / mRNA length | 278,993,619 | 170,627,661 | 90,669,335 | 103,342,350 | 106,564,991 | 122,281,322 |
| Total exon length | 49,834,014 | 29,734,201 | 22,409,584 | 22,298,859 | 22,288,118 | 25,630,404 |
| Total CDS pieces length | 33,362,898 | 29,578,314 | 21,700,488 | 21,984,588 | 22,028,601 | 25,377,594 |
| Total 5' UTR length | 1,942,114 | 122,107 | 80,708 | 73,260 | 79,275 | 114,997 |
| Total 3' UTR length | 14,529,002 | 33,780 | 628,388 | 241,011 | 180,242 | 137,813 |
| Total intron length | 229,356,734 | 141,056,991 | 68,365,309 | 81,150,437 | 84,383,113 | 96,782,096 |
| Mean gene / mRNA length | 13,832 | 7,665 | 3,808 | 4,164 | 4,371 | 5,353 |
| Mean total CDS length in genes | 1,654 | 1,328 | 911 | 885 | 903 | 1,111 |
| Mean exon length | 229 | 160 | 173 | 169 | 170 | 166 |
| Mean 5' UTR length | 220 | 37 | 56 | 52 | 56 | 33 |
| Mean 3' UTR length | 1,465 | 375 | 834 | 554 | 541 | 443 |
| Mean CDS pieces length | 162 | 159 | 172 | 168 | 170 | 165 |
| Mean intron length | 1,163 | 862 | 647 | 758 | 794 | 737 |
| Proportion of repeat elements in genome (%) | 30.45 | 30.73 | 29.66 | 24.83 | 39.12 | 34 |
| Proportion of exons in genome (%) | 8.78 | 5.16 | 4.00 | 4.19 | 3.12 | 4.03 |
| Proportion of introns in genome (%) | 40.39 | 24.50 | 12.19 | 15.26 | 11.82 | 15.21 |
| Proportion of genes in genome (%) | 49.16 | 29.66 | 16.19 | 19.45 | 14.94 | 19.23 |

Note: Gene = Exons + Introns. UTR = untranslated region. CDS = coding sequences = exon - UTR regions.

In all assemblies, the largest classified repeat element proportion was always DNA transposons (6.92%–14.85%), followed by long interspersed nuclear element (LINE) retrotransposons (3.26%–5.83%) (Figure 4, Supplementary Table 2). The proportion of repeat elements that were not classified in the Dfam/RepBase databases was 10.53%–13.49%. Long terminal repeat (LTR) retrotransposons (1.29%–1.82%), simple interspersed nuclear element (SINE, 0.45%–0.50%) and simple repeats (1.18%–1.71%) represented a smaller fraction in comparison.

**Figure 4.**
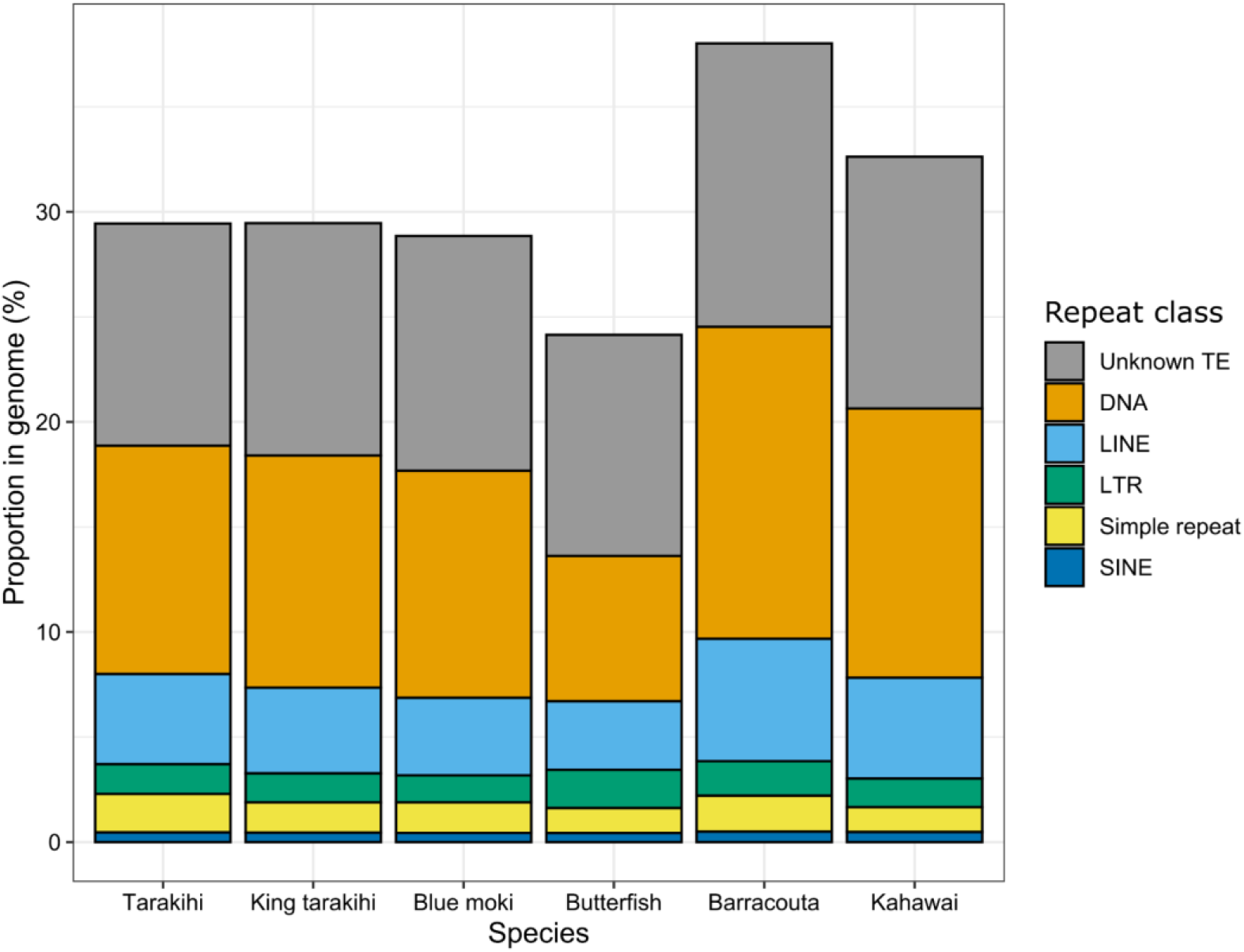
Proportions of the main classes of repeat elements in the genomes. Repeat elements are either simple repeats or transposable elements (TE), which include DNA transposons (DNA), long terminal repeat (LTR) retrotransposons, and non-LTR retrotransposons (long and short interspersed nuclear elements, LINE and SINE).

To assess the proportion of repeat elements in more detail, the ten most frequent repeat families were identified for each species (Figure 5, Supplementary Figure 1). The ten most frequent repeat families were always the same in tarakihi, king tarakihi, blue moki, barracouta and kahawai: they were unclassified elements, simple repeats, DNA hAT-Ac, LINE L2, DNA hAT-Tip100, DNA TcMar-Tc1, LINE Rex-Babar, unknown DNA elements, DNA PIF-Harbinger, and DNA hAT-Charlie (not always in the same order). For butterfish, the ten most frequent repeat families were almost all the same as the other species, with two exceptions: LTR Ngaro and rolling-circle helitrons were part of the ten most frequent repeat elements in that species, instead of the unknown DNA elements and DNA hAT-Tip100.

**Figure 5.**
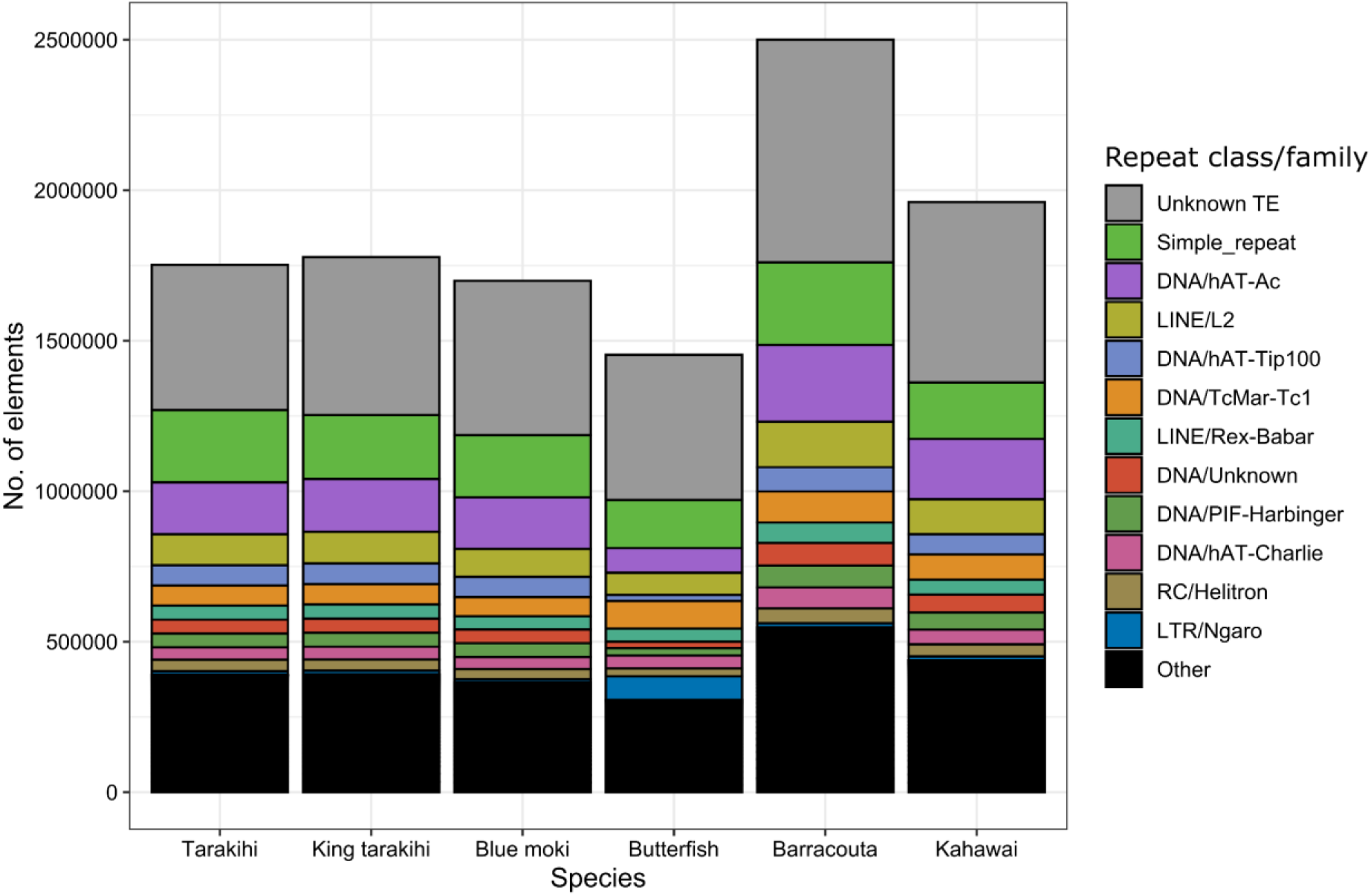
Proportions of the most represented families of repeat elements in the genomes. Repeat elements families are sorted vertically based on their abundance in tarakihi. “Other” includes all the families that are not in the top ten of the most abundant RE in at least one species. See Supplementary Figure 1 for more details on the repeat element included in this group.

The number of genes detected by the annotation pipeline in the five new genome assemblies ranged from 22,258 (king tarakihi) to 24,816 (butterfish), which is higher than the 20,169 genes found in tarakihi (Table 2). The mean number of exons per gene ranged from five (blue moki, butterfish, and barracouta) to eight (king tarakihi), both lower mean values when compared to the eleven found for tarakihi (Table 2). Many of the other gene feature metrics were comparable across species, with the largest deviations seen for tarakihi and king tarakihi (Table 2), see Discussion.

### Genomic comparative analysis

Gene family clustering of the 14 fish species assigned 96.3% of all the genes in orthogroups. A total of 19,874 orthogroups were found, among which 1,481 were single-copy orthogroups that were used for phylogenetic reconstruction (Figure 6). There were 11 orthogroups only specific to tarakihi, while the number for the five other New Zealand species varied from 4 to 71 (Supplementary Table 3). Genes contained only in the tarakihi-specific orthogroups were: coxsackievirus and adenovirus receptor (four copies), endonuclease-reverse transcriptase (three copies), poly(rC)-binding protein, small G protein signalling modulator 1, paired box protein Pax-7, glucagon-1-like, sulfhydryl oxidase 2 (two copies) and a few unannotated genes. Tarakihi, spotted gar, and green spotted puffer were the only three species with a negative average gene expansion, meaning they displayed a net loss of genes per gene family (Supplementary Table 4).

**Figure 6.**
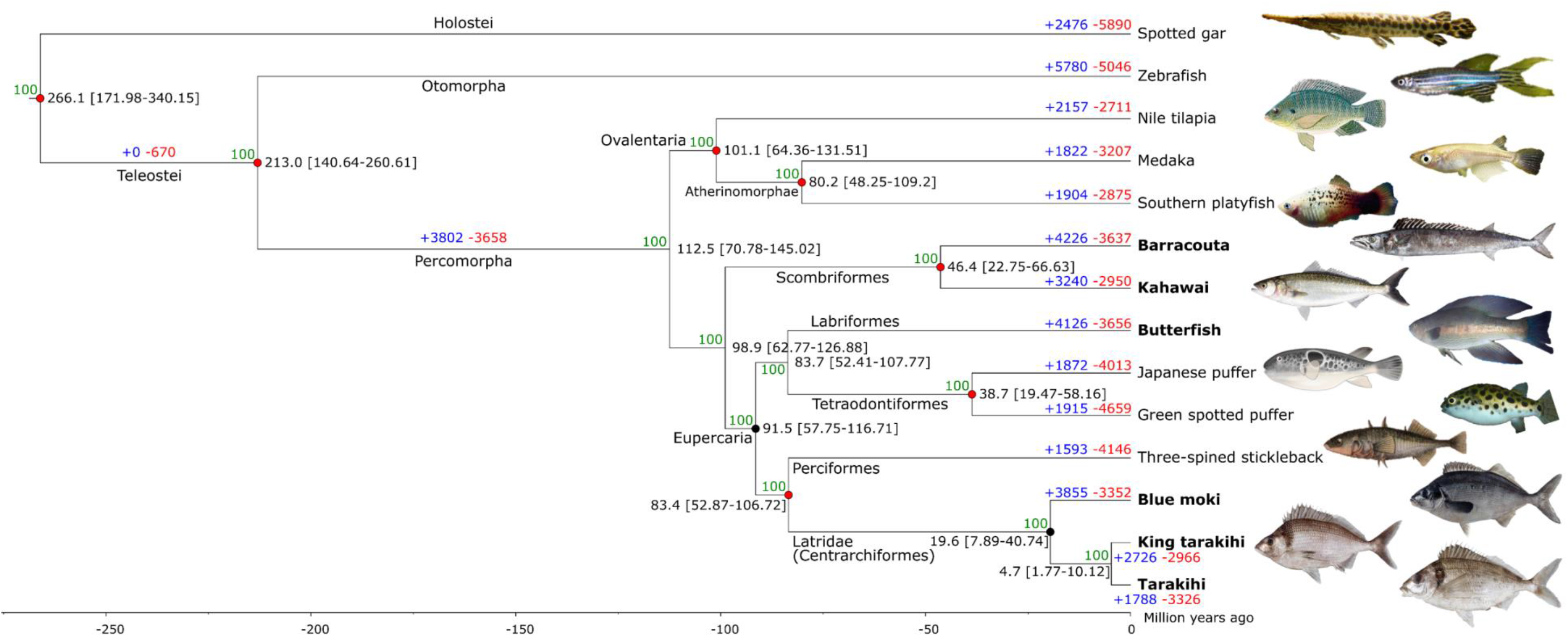
Phylogenomic tree of the six New Zealand fish species (in bold) and eight other fish species. A dot on a node indicates that the clade is monophyletic *sensu* Betancur-R et al. (2017). The red dots in particular have been used for node age calibration with TimeTree. Black numbers indicate the estimated divergence time with 95% confidence intervals. Blue and red numbers represent respectively the expansion and the contraction of gene families along the phylogeny. Green numbers are the posterior probabilities in percentage and indicate the support values of the nodes. See the “pictures credits” section below for fish pictures attributions.

Overall the phylogenetic tree (Figure 6) was consistent with recently reported molecular phylogenies of Percomorpha (Betancur-R et al., 2017; Sanciangco et al., 2016; Shou & Han, 2021). All nodes were strongly supported (posterior probability = 100%) and orders were retrieved as expected. The Latridae family (which include tarakihi, king tarakihi, and blue moki), representing the order Centrarchiformes, was placed as a sister clade of “true” Perciformes (stickleback), which is consistent with Betancur-R et al. (2017). There was one difference with the phylogenies from Betancur-R et al. (2017), Sanciangco et al. (2016) and Shou & Han (2021): the positions of the Pelagiaria (Scombriformes: Barracouta and Kahawai) and the Ovalentaria clades (which include Cyprinodontiformes, Beloniformes, and Cichliformes) are swapped. In Eupercaria, the Tetraodontiformes (pufferfishes) were placed as a sister clade of Labriformes (wrasses: butterfish) in our phylogeny, which is consistent with Shou & Han (2021) but not with Betancur-R et al. (2017) and Sanciangco et al. (2016). These studies placed pufferfishes as a sister clade of Perciformes + Centrarchiformes instead. This is of particular interest because all the species newly assembled here belong to the clade Percomorpha (Percomorphaceae *sensu* Betancur-R et al. (2017)). This taxonomic clade is historically considered a “bush at the top” of the fish tree of life that includes around 55% of extant bony fish species (Sanciangco et al., 2016). Consequently, both the phylogenetic inter-relationships between the large clades cited above (Ovalentaria, Pelagiaria (Scombriformes), and Euparcaria) and the position of Tetraodontiformes in Eupercaria are still contentious, with nodes supports being relatively low even in recent studies (Betancur-R et al., 2017; Sanciangco et al., 2016).

According to the present phylogeny, the radiation of Percomorpha has occurred c. 112 million years ago, in the mid-Cretaceous (Figure 6). The Centrarchiformes (tarakihi, king tarakihi, and blue moki) have then diverged from the true Perciformes c. 83 million years ago, in the late Cretaceous. Tarakihi and king tarakihi are estimated to have diverged 4.7 million years ago. While still relatively recent, this is older than the minimum time since divergence of 0.3–0.8 million years estimated with nucleotide divergence rate of the mitochondrial control region (Papa, Halliwell, et al., 2021). The confidence interval is, however, comparatively large (95% CI = 1.77–10.12 MYA), and the “true” time since divergence could be situated somewhere in the lower end of that interval.

A total of 65 genes in the tarakihi genome showed evidence consistent with positive selection (Supplementary Table 5). These 65 genes under selection were associated with 295 GO terms for biological processes. The GO terms that appeared the most often (i.e. more than two times) included terms related to function: ATP binding, metal ion binding, regulation of transcription by RNA polymerase II, integral component of membrane, RNA polymerase II cis-regulatory region sequence-specific DNA binding, RNA binding, zinc ion binding, DNA-binding transcription factor activity (RNA polymerase II-specific), GTPase activator activity, signal transduction, microtubule binding, hydrolase activity, protein dimerization activity, and cell division (respectively). They also included terms related to location: cytoplasm, nucleus, cytosol, endosome, Golgi apparatus (respectively) (Figure 7).

**Figure 7.**
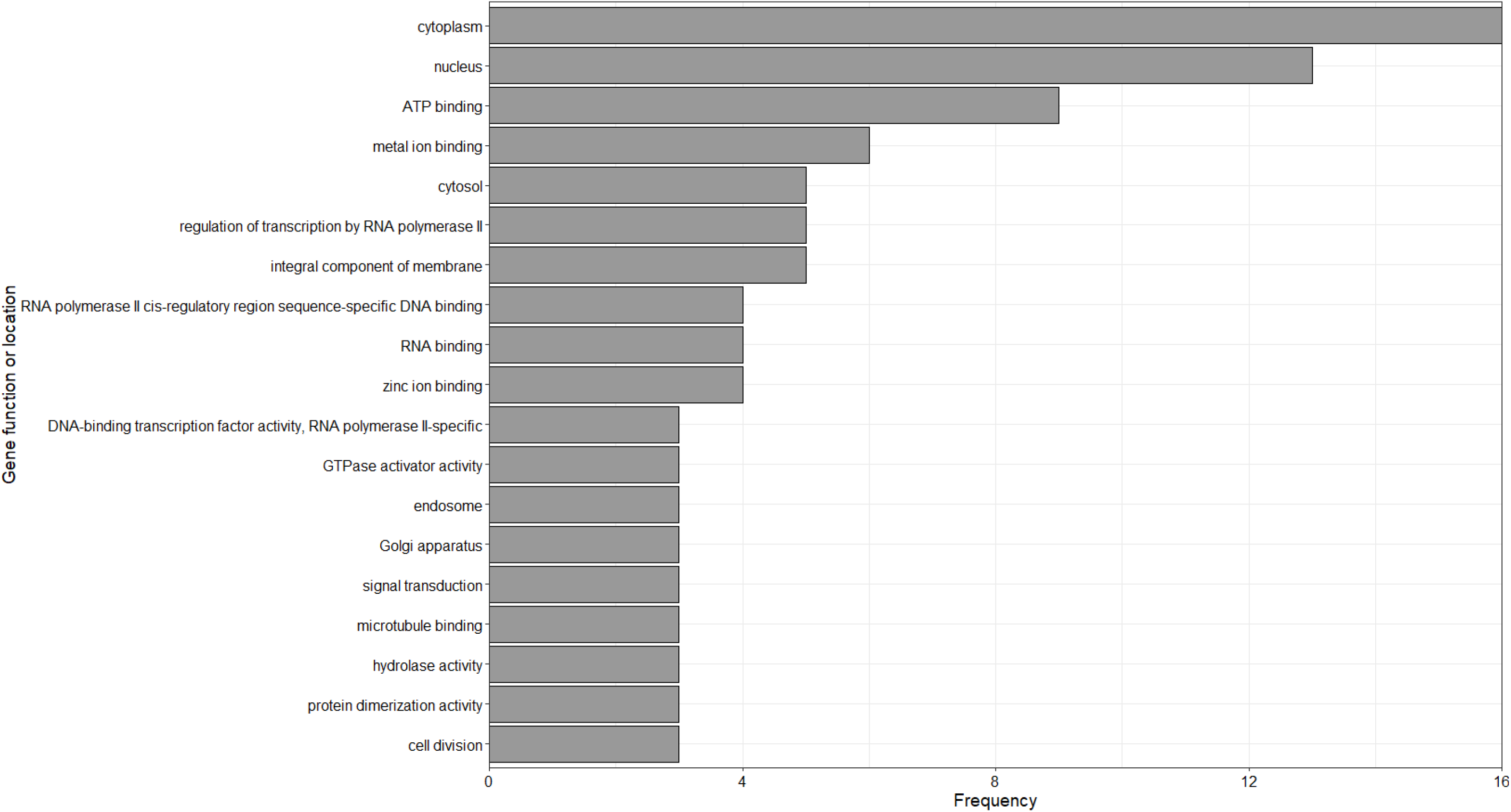
Frequency of gene functions and locations according to GO terms for genes under positive selection in tarakihi. Only terms associated with three genes or more are shown.

A total of 209 genes showed evidence of positive selection in the Latridae lineage, which included tarakihi, king tarakihi and blue moki (Supplementary Table 6). They included almost all of the genes detected in tarakihi except for six: cell division cycle 25B, high mobility group 20A, nuclear receptor subfamily 2 group C member 2, peter pan homolog, ribosomal protein L27, and zgc:85936 were positively selected in the tarakihi only (Supplementary Table 5). The most frequent GO terms for selected genes functions in Latridae were integral component of membrane, ATP binding, metal ion binding, RNA binding, regulation of transcription by RNA polymerase II, zinc ion binding, RNA polymerase II cis-regulatory region sequence-specific DNA binding, DNA-binding transcription factor activity (RNA polymerase II-specific), respectively. The most frequent location terms were cytoplasm, nucleus, cytosol, plasma membrane, and membrane, respectively (Figure 8).

**Figure 8.**
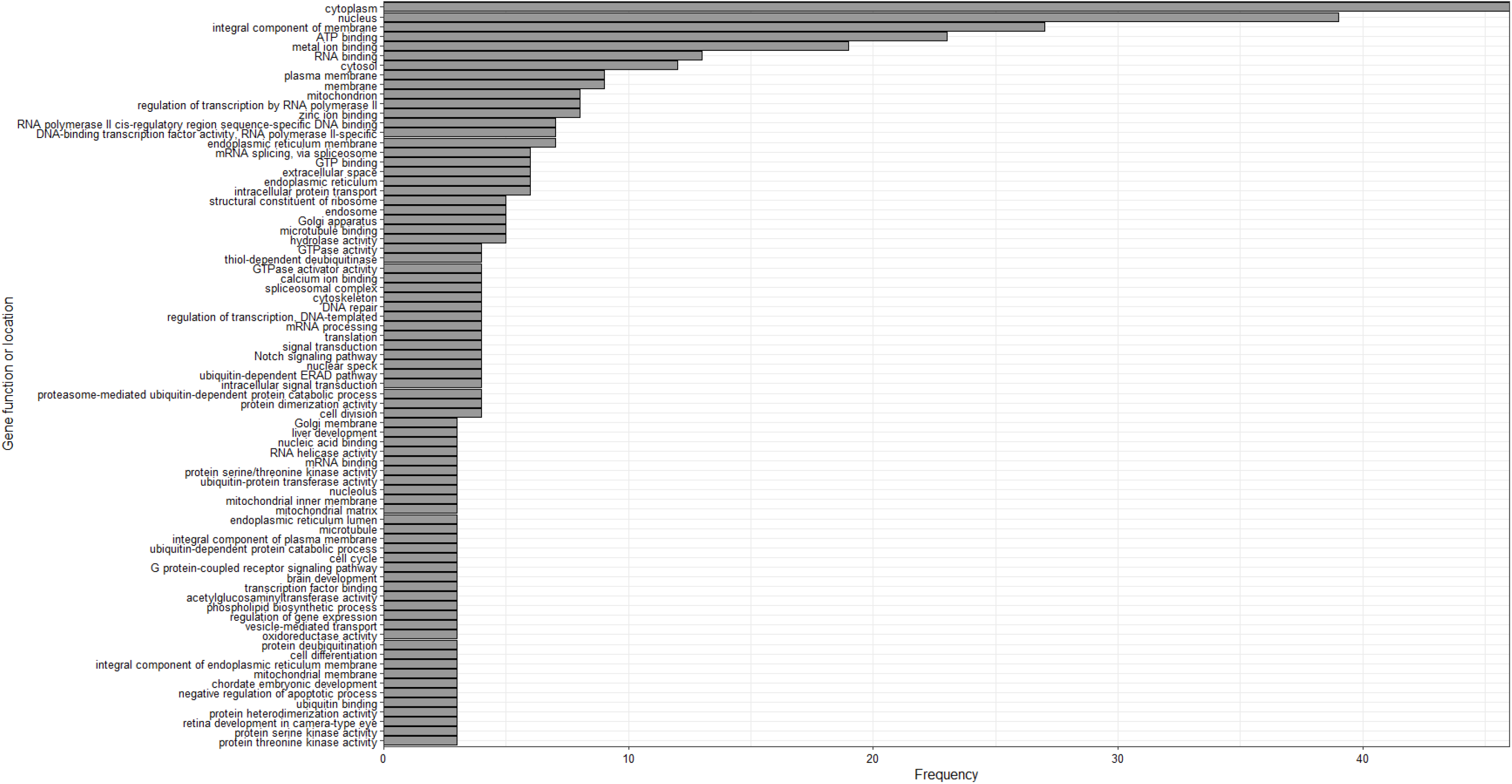
Frequency of gene functions and locations according to GO terms for genes under positive selection in Latridae. Only terms associated with three genes or more are shown.

## Discussion

This study provided the first genome assemblies for five New Zealand fish species. This contribution fills an important gap in the genomic resources for the South Pacific ichthyofauna. The analyses of the genomes enabled a range of genomic features to be identified and described, in particular the diversity of repeat elements and the most represented repeat families. The phylogenetic analysis of the combined data set of 14 species provided insights into the evolutionary history of fish and the effects of selection on fish genomes.

### Repetitive elements in fish genomes

DNA transposons represented the highest proportion of repeat elements in all six genome assemblies, making up more than 10% of the total genome sequence in all species except butterfish (Figure 4, Supplementary Table 2). DNA transposons differ from retrotransposons in that they do not rely on an RNA intermediate for moving within the genome (Sotero-Caio et al., 2017; Wicker et al., 2007). This finding is consistent with other comparative analyses that reported DNA transposons as the most abundant class of repeat elements in most teleost fish species (Brawand et al., 2014; Chalopin et al., 2015; Gao et al., 2016; Shao et al., 2019). While all six species investigated here belong to the large Percomorpha clade, one of the most comprehensive studies in terms of species numbers (35 Actinopterygii from 14 orders) showed that while DNA transposons are often the dominant RE class in ray-finned fishes, including Percomorpha, they can in some cases be outnumbered by LINE or LTR retrotransposons (Shao et al., 2019). This was the case for the three-spined stickleback (a “true” Perciform, which is the sister taxon of Centrarchiformes that include tarakihi, king tarakihi and blue moki) and other Percomorpha with reduced genome size like Japanese puffer and green spotted puffer. Also consistent with our results, short interspersed nuclear element (SINE) transposons usually represent a much smaller fraction of the genome in fishes (Shao et al., 2019) compared to e.g. mammals (Sotero-Caio et al., 2017).

Observing the frequencies of RE at the intra-class level showed that all six species, except butterfish, shared the same most common TEs (Figure 5). These were comprised of three TE families from the hAT superfamily of DNA transposons (Ac, Tip100, and Charlie), two other DNA transposons superfamilies (Tc1/Mariner, mostly represented by the Tc1 family, and PIF/Harbinger), and two superfamilies of LINE retrotransposons (L2 and Rex/Babar). Albeit not always in the exact same order, the relative proportion of each of these families were similar in tarakihi, king tarakihi, blue moki, barracouta and kahawai (Figure 5). hAT, Tc1/mariner and PIF/Harbinger are “cut-and-paste” DNA transposons. “Cut-and-paste” DNA TEs move among genomic locations via excision and insertion using a transposase (Y. W. Yuan & Wessler, 2011). L2 and Rex/Babar are long interspersed nuclear elements (LINEs). LINEs are retrotransposons that contain a reverse transcriptase gene but lack long terminal repeats (Finnegan, 2012). These results are consistent with a former extensive comparative study that found that Tc1/Mariner, hAT, and L2 superfamilies are among the most important TE in fish genomes both in terms of genome proportion and representation across taxa (Shao et al., 2019), although they also included L1 and Gypsy elements in that list. Interestingly, butterfish was the only one of the six species that showed a different trend in most represented RE elements, with an over-representation of Ngaro elements (a type of Long Terminal Repeats retrotransposon) compared to e.g. hat-Tip100 DNA transposons (Figure 5). While Ngaro elements do not seem to be especially dominant in fishes overall in comparison to other LTRs like e.g. Gypsy elements (Shao et al., 2019), they were found to differ highly in terms of abundance and diversity among teleost lineages (Gao et al., 2016). Indeed this trend is also observed in these six fish species. Finally, while the helitron elements were among the ten most common TE families in butterfish only, they were not especially more abundant in comparison with the five other species, where they also represent a sizeable proportion of all TE (Figure 5). Helitrons are a specific kind of DNA transposon that do not move via “cut-and-paste” but rather use a “rolling-circle” replication mechanism (Wicker et al., 2007).

A strong and significant relationship between the proportion of repeat elements and the genome size was found (Figure 3). This observation was especially robust since it did not depend on the quality of the genome assembly (Figure 9) (see below). This result provides additional evidence for the strong relationship between the genome size and the abundance of REs that has already been observed in a broad number of fish taxa (Gao et al., 2016; Shao et al., 2019; Z. Yuan et al., 2018). Current evidence suggests that the genome size of ray-finned fishes is more dependent on repeat elements in general than for tetrapods (Chalopin et al., 2015; Sotero-Caio et al., 2017). Specifically, it has been suggested that DNA transposons could be significant drivers of the genome size differentiation in teleost fish, whereas LTR and non-LTR retrotransposons seem to dominate the genome expansion of most reptiles and mammals (Gao et al., 2016).

**Figure 9.**
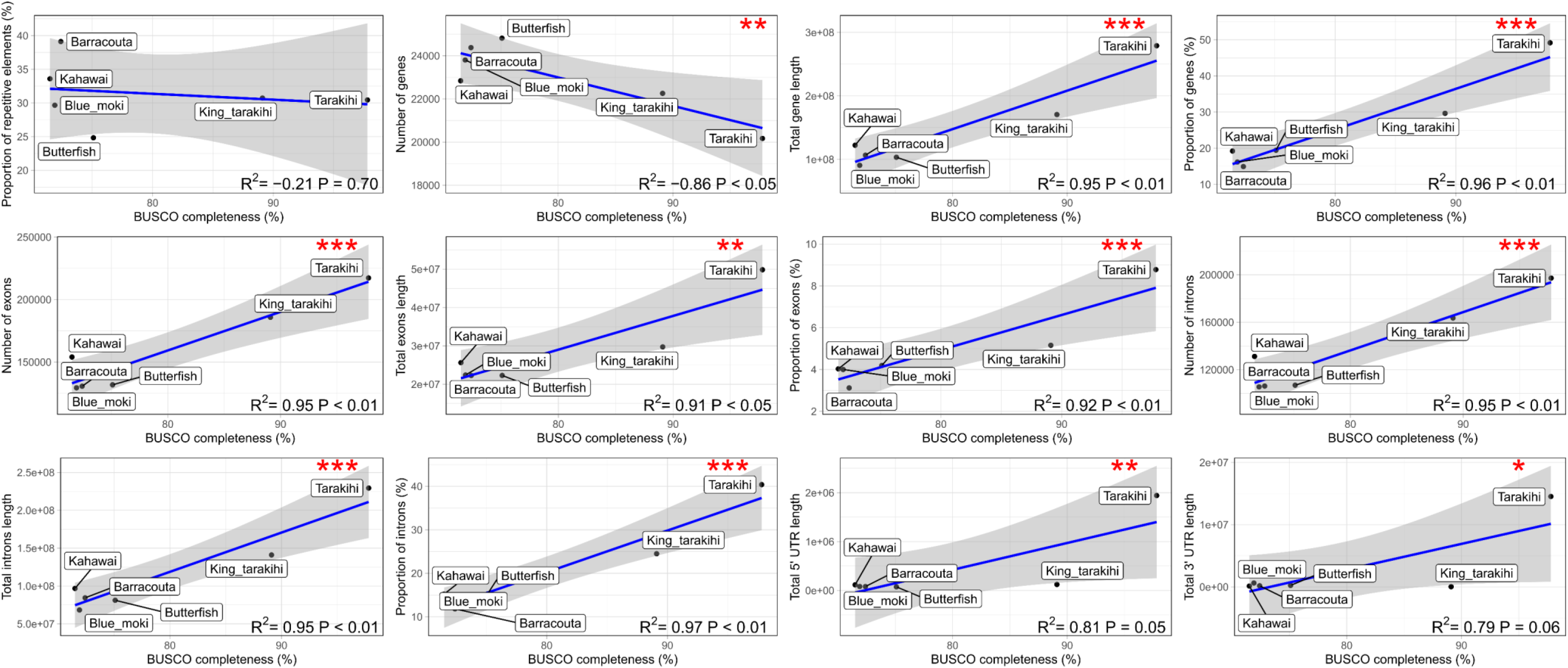
Correlation between BUSCO completeness and proportion, number and length of genomic features annotated in the six genome assemblies, with corresponding Pearson correlation coefficient (*R*^2^) and p-value (*P*). Grey area is the 95% confidence interval. Red asterisks indicate significance, with *P* ≤ 0.1 (*), 0.05 (**), and 0.01 (***).

### Impact of genome size and fragmentation on genome annotation quality

Genomic features were detected through the annotation pipeline in the newly assembled fish genomes (Table 2). One of the goals of this study was to explore the genomic features (i.e. number and proportion of both repeat elements and genes and their constituents) in more detail to infer trends across species. However, it was necessary to test for potential biases induced by the genome assembly quality beforehand for the interpretation of the results to be robust. We found that all genomic features explored, with the notable exception of the proportion of repeat elements, correlated strongly with genome completeness (Figure 9) as opposed to e.g. genome size (Figure 10). Higher fragmentation (i.e. lower contiguity) of the genome assembly reduces the length of genes detected and reduces both the number and length of the genic components, including exons, introns and UTR regions (Figure 9). It is possible that a higher fragmentation creates a bias toward the detection of smaller genomic features. However, the number of genes decreased when the assembly contiguity was higher. This could be due to several reasons. First, the higher fragmentation of the genomes might increase the quantity of erroneously unmerged gene duplicates. This result could also be due to differences in the annotation pipeline. The tarakihi and the king tarakihi genomes, which also happen to be the two most highly continuous assemblies, have been annotated using full-length RNA reads (Iso-Seq) instead of short RNA-seq data. The RNA reads used for the tarakihi were also the only ones coming from the same species as opposed to a close species: this led to some slight changes in the annotation pipeline parameters for the five other species compared to tarakihi, which might have influenced the gene discovery process by making it less stringent.

**Figure 10.**
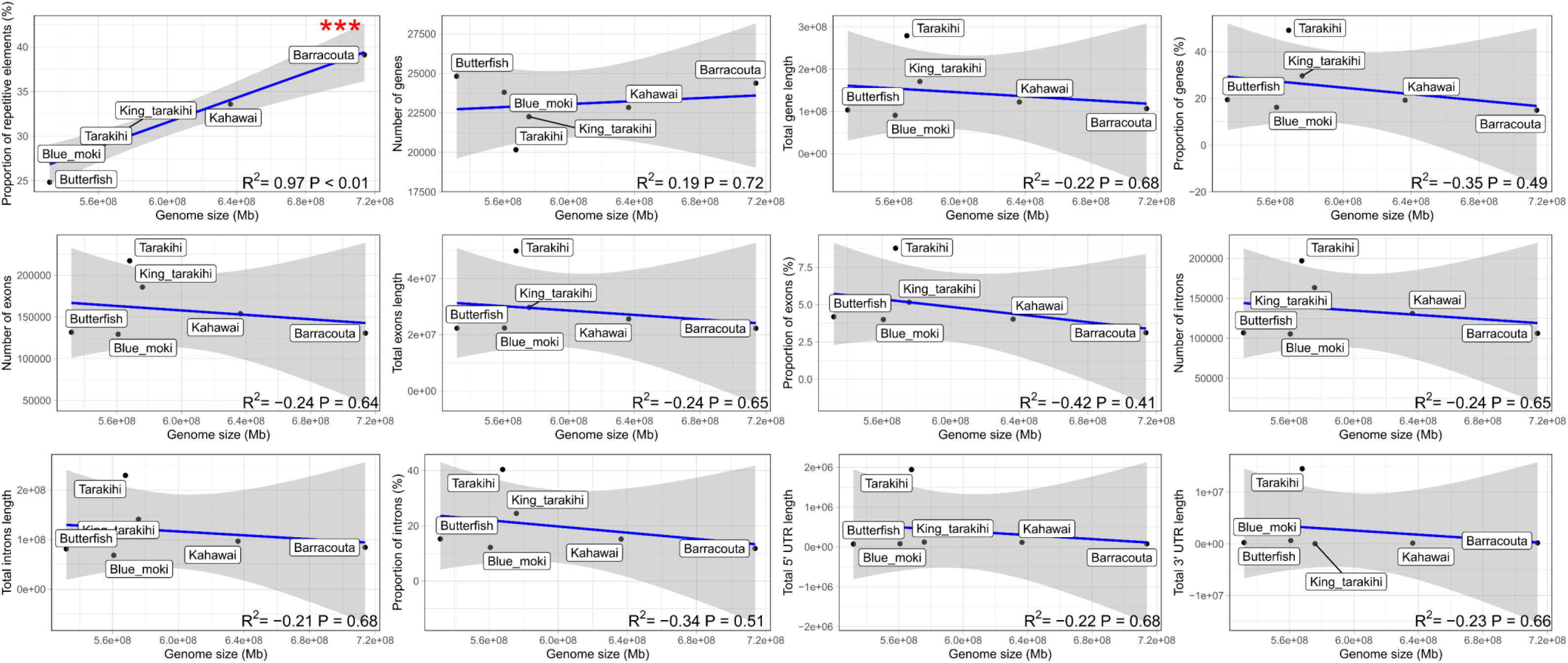
Correlation between genome size and proportion, number and length of genomic features annotated in the six genome assemblies, with corresponding Pearson correlation coefficient (R2) and p-value (P). Grey area is the 95% confidence interval. Red asterisks indicate significance, with P ≤ 0.1 (*), 0.05 (**), and 0.01 (***)

### Positive selection in ATP binding genes

The most frequent GO term for function in tarakihi was “ATP binding”. The term was associated with nine of the positively selected genes detected in tarakihi, which means that more than 10% of its 65 positively selected genes are related to that function. It was also the most frequent function of the selected genes for Latridae just after “integral component of membrane”, although in most cases the two functions were related to the same genes. Genes in the ATP binding pathway provide energy for the active transport of ions across cell membranes (Melkikh & Seleznev, 2012). Genes in the ATP binding pathway were the most common genes found to be under positive selection in a former study based on molecular evolution rates of four model fish species (Steinke et al., 2006) and were also found to be positively selected in the largemouth bass (Sun et al., 2021), another fish of the order Centrarchiforme. Mutations in genes involved in ATP binding, especially the ATP-Binding Cassette transporter genes, are known to be associated with a wide variety of pathologic disorders (Dean & Annilo, 2005). Taken together, these results may imply that ATP binding genes are submitted to both strong purifying selection against deleterious mutations and positive selection favouring beneficial mutations. Further research will be needed to assess if the propensity of these genes to fix novel mutations is related to the speciation process, and if it is a driver or a result of this process.

## Conclusion

The genome sequence of five New Zealand bony fish species, six including the tarakihi, provided several results relevant to shed light on the genomic features associated with the evolution of these species. In addition to their utility in a comparative genomic study, the genome assemblies can be used as references for population genomics studies. While it always depends on the question addressed, if the resources are available, the aim for these types of applications should ideally be to obtain a sex-specific, phased, chromosome-level genome with transcriptome-informed annotation. In these cases, long-read DNA sequencing (e.g. PacBio Hi-Fi reads, ONT reads), supplemented with scaffolding data (e.g. Hi-C), isoform data (e.g. Iso-Seq) and sometimes short high-quality reads (e.g. Illumina short reads, DNBSeq) should be used. However, given the decreasing costs and fewer sampling constraints, the present pipeline could be an efficient way to easily assemble the genomes of more fish species, especially the ones for which funding is less available because of lower commercial importance.

The study also provided a molecular phylogeny based on whole genomes for the contentious clade Percomorphaceae with strong node supports. Adding more species to the tree, both with newly assembled genomes and from genomic data already available will surely provide further insight into the complicated evolutionary history of this clade that represents a considerable portion of living vertebrates. Finally, we found evidence of positive selection on genes involved in the ATP binding pathway, consistent with other studies of fish. Further studies of the molecular and biological processes involved in these pathways and their putative association with environmental factors will be needed to better apprehend the benefits of maintained novel mutations in these genes.

## Acknowledgements

This research was supported by the programme “Juvenile fish habitat bottlenecks” (CO1X1618), funded by the New Zealand Ministry of Business, Innovation, and Employment (MBIE) Endeavour Fund. We are grateful to Dive Wellington recreational fishermen, Nick Johnston, and Moana NZ for providing the fish specimens used in this study. All fish pictures used in the phylogenomic tree are either courtesy of the Museum of New Zealand Te Papa Tongarewa or derived from work in the public domain, except for the following: *Lepisosteus oculatus* photo by Brian Gratwicke: https://commons.wikimedia.org/wiki/File:Lepisosteus_oculatus1.jpg / [CC-BY-2.5](https://creativecommons.org/licenses/by/2.5/deed.en), *Oryzias latipes* photo by NOZO: https://upload.wikimedia.org/wikipedia/commons/4/49/Nihonmedaka.jpg / [CC-BY-3.0](https://creativecommons.org/licenses/by/3.0/deed.en), *Xiphophorus maculatus* photo by vxixiv: https://www.flickr.com/photos/21630815@N06/3406554064 / [CC-BY-2.0](https://creativecommons.org/licenses/by/2.0/deed.en), *Takifugu rubripes* photo by DataBase Center for Life Science (DBCLS): https://doi.org/10.7875/togopic.2012.10 / [CC-BY-4.0](https://creativecommons.org/licenses/by/4.0/deed.en).

## Data Accessibility

All data generated for this study can be accessed upon request on the Genomics Aotearoa 647 repository (https://repo.data.nesi.org.nz/) under project name “tarakihi genomics”. All bash and R scripts used for this study are available on GitHub in the following repository: https://github.com/yvanpapa/comparative_genomics_NZ_fish.

## CRediT authorship contribution statement

YP: Conceptualization, Methodology, Software, Validation, Formal analysis, Investigation, Resources, Data Curation, Writing - Original Draft, Writing - Review & Editing, Visualization. MW & MM: Resources, Writing - Review & Editing, Supervision, Funding acquisition. PR: Conceptualization, Resources, Writing - Review & Editing, Supervision, Project administration, Funding acquisition.

## Notes

### Competing Interest Statement

The authors have declared no competing interest.

