## Supplementary_Material for "Comparative genomics of tarakihi (*Nemadactylus macropterus*) and five New Zealand fish species: assembly contiguity affects the identification of genic features but not transposable elements"

### Supplementary Tables

Supplementary Table 1. Summary of number of reads and bases obtained at several steps of the quality filtering pipelines.

| | | Raw reads | Quality-filtered reads | Uncontaminated reads | Mitochondrial reads (%) | Final Illumina PE reads |
|---|---|---|---|---|---|---|
| King tarakihi | No. reads | 188,470,220 | 170,505,602 | 156,465,038 | 0.14997344 | 156,230,382 |
| | No. bases | 28,270,533,000 | 25,575,840,300 | 23,469,755,700 | 0.14997344 | 23,434,557,300 |
| Barracouta | No. reads | 205,858,992 | 184,329,218 | 158,680,852 | 0.034627997 | 158,625,904 |
| | No. bases | 30,878,848,800 | 27,649,382,700 | 23,802,127,800 | 0.034627997 | 23,793,885,600 |
| Blue moki | No. reads | 189,522,608 | 174,855,726 | 160,748,294 | 0.151186674 | 160,505,264 |
| | No. bases | 28,428,391,200 | 26,228,358,900 | 24,112,244,100 | 0.151186674 | 24,075,789,600 |
| Butterfish | No. reads | 219,182,928 | 198,607,764 | 186,745,068 | 0.983231322 | 184,908,932 |
| | No. bases | 32,877,439,200 | 29,791,164,600 | 28,011,760,200 | 0.983231322 | 27,736,339,800 |
| Kahawai | No. reads | 232,301,418 | 211,988,224 | 197,198,048 | 0.201719035 | 196,800,262 |
| | No. bases | 34,845,212,700 | 31,798,233,600 | 29,579,707,200 | 0.201719035 | 29,520,039,300 |

Supplementary Table 2. Main classes and proportions of repeat elements detected in the five new assembled genomes

| | King tarakihi | Barracouta | Blue moki | Butterfish | Kahawai |
|---|---|---|---|---|---|
| RNA-mediated class I transposons (retrotransposons) | | | | | |
| SINEs | 33677 0.45% | 40399 0.50% | 31713 0.44% | 32533 0.43% | 36814 0.48% |
| Penelope | 8207 0.14% | 14509 0.26% | 8222 0.14% | 9252 0.14% | 11805 0.22% |
| LINEs | 217366 4.08% | 303643 5.83% | 198643 3.69% | 180766 3.26% | 238505 4.80% |
| LTR elements | 76131 1.38% | 96989 1.64% | 71646 1.29% | 110681 1.82% | 74939 1.36% |
| RNA-independent class II DNA transposons | | | | | |
| DNA transposons | 591041 11.05% | 874958 14.85% | 568932 10.81% | 388300 6.92% | 692761 12.82% |
| Rolling-circles | 34869 0.46% | 46474 0.64% | 32087 0.42% | 24792 0.36% | 37629 0.60% |
| Unclassified transposable elements | 502186 11.06% | 701196 13.49% | 490965 11.17% | 466526 10.53% | 571345 11.98% |
| Small RNA | 8989 0.11% | 12665 0.14% | 8695 0.11% | 7875 0.10% | 10716 0.13% |
| Satellites | 8263 0.51% | 9219 0.20% | 5925 0.11% | 4487 0.08% | 6646 0.12% |
| Simple repeat | 211399 1.44% | 273196 1.71% | 205792 1.45% | 159110 1.19% | 186620 1.18% |
| Low complexity | 27126 0.25% | 30005 0.21% | 23797 0.23% | 20738 0.20% | 26541 0.21% |
| Total REs | 30.73% | 39.12% | 29.66% | 24.83% | 33.59% |

Supplementary Table 3. Statistics of orthogroups per species

| | Tarakihi | King tarakihi | Barracouta | Blue moki | Butterfish | Kahawai | Zebrafish | Stickleback | Spotted gar | Nile tilapia | Medaka | Takifugu | Tetraodon | Platyfish |
|---|---|---|---|---|---|---|---|---|---|---|---|---|---|---|
| No. of genes | 20169 | 22258 | 24378 | 23804 | 24816 | 22840 | 30313 | 20787 | 18341 | 28189 | 23622 | 21411 | 19602 | 23774 |
| No. of genes in orthogroups | 19745 | 21479 | 22627 | 22369 | 22697 | 22163 | 29333 | 20112 | 17908 | 27621 | 22889 | 20845 | 19342 | 23226 |
| No. of unassigned genes | 424 | 779 | 1751 | 1435 | 2119 | 677 | 980 | 675 | 433 | 568 | 733 | 566 | 260 | 548 |
| % of genes in orthogroups | 97.9 | 96.5 | 92.8 | 94 | 91.5 | 97 | 96.8 | 96.8 | 97.6 | 98 | 96.9 | 97.4 | 98.7 | 97.7 |
| % of unassigned genes | 2.1 | 3.5 | 7.2 | 6 | 8.5 | 3 | 3.2 | 3.2 | 2.4 | 2 | 3.1 | 2.6 | 1.3 | 2.3 |
| No. of orthogroups containing species | 14126 | 14430 | 14458 | 14644 | 14404 | 14355 | 14053 | 13112 | 13060 | 14441 | 14037 | 13272 | 12559 | 14247 |
| % of orthogroups containing species | 71.1 | 72.6 | 72.7 | 73.7 | 72.5 | 72.2 | 70.7 | 66 | 65.7 | 72.7 | 70.6 | 66.8 | 63.2 | 71.7 |
| No. of species-specific orthogroups | 11 | 4 | 52 | 39 | 71 | 12 | 318 | 29 | 73 | 207 | 91 | 66 | 33 | 68 |
| No. of genes in species-specific orthogroups | 25 | 8 | 112 | 81 | 149 | 26 | 2463 | 278 | 352 | 1701 | 772 | 256 | 116 | 346 |
| % of genes in species-specific orthogroups | 0.1 | 0 | 0.5 | 0.3 | 0.6 | 0.1 | 8.1 | 1.3 | 1.9 | 6 | 3.3 | 1.2 | 0.6 | 1.5 |

Supplementary Table 4. Statistics of gene family expansions and contractions

| Species | Expanded fams | Genes gained | genes/expansion | Contracted fams | Genes lost | genes/contraction | No change | Avg. Expansion |
|---|---|---|---|---|---|---|---|---|
| Barracouta | 4226 (4226) | 7008 | 1.66 | 3637 (3637) | 4240 | 1.17 | 10930 | 0.147289 |
| Blue moki | 3855 (3855) | 6428 | 1.67 | 3352 (3352) | 3885 | 1.16 | 11586 | 0.135316 |
| Butterfish | 4126 (4126) | 7058 | 1.71 | 3656 (3656) | 4256 | 1.16 | 11011 | 0.149098 |
| Kahawai | 3240 (3240) | 5804 | 1.79 | 2950 (2950) | 3413 | 1.16 | 12603 | 0.127228 |
| King tarakihi | 2726 (2726) | 5119 | 1.88 | 2966 (2966) | 3394 | 1.14 | 13101 | 0.0917895 |
| Medaka | 1822 (1822) | 5799 | 3.18 | 3207 (3207) | 3492 | 1.09 | 13764 | 0.122758 |
| Nile tilapia | 2157 (1672) | 8586 | 3.98 | 2711 (188) | 2935 | 1.08 | 13925 | 0.300697 |
| Platyfish | 1904 (1904) | 6172 | 3.24 | 2875 (2875) | 3111 | 1.08 | 14014 | 0.16288 |
| Spotted gar | 2476 (787) | 4427 | 1.79 | 5890 (2) | 5898 | 1 | 10427 | -0.0782738 |
| Stickleback | 1593 (1593) | 4695 | 2.95 | 4146 (4146) | 4610 | 1.11 | 13054 | 0.00452296 |
| Takifugu | 1872 (1872) | 5241 | 2.8 | 4013 (4013) | 4511 | 1.12 | 12908 | 0.0388443 |
| Tarakihi | 1788 (1788) | 3757 | 2.1 | 3326 (3326) | 3783 | 1.14 | 13679 | -0.00138349 |
| Tetraodon | 1915 (1915) | 4684 | 2.45 | 4659 (4659) | 5225 | 1.12 | 12219 | -0.0287873 |
| Zebrafish | 5780 (2419) | 13130 | 2.27 | 5046 (0) | 5046 | 1 | 7967 | 0.43016 |

Supplementary Table 5. Genes positively selected in the tarakihi [Continues next page]

| Gene | Protein name |
|---|---|
| abhd11 | abhydrolase domain containing 11 |
| acmsd | aminocarboxymuconate semialdehyde decarboxylase |
| apex1 | APEX nuclease (multifunctional DNA repair enzyme) 1 |
| arhgap24 | Rho GTPase activating protein 24 |
| arl13a | ADP-ribosylation factor-like 13A |
| cabin1 | calcineurin binding protein 1 |
| cdc25b* | cell division cycle 25B |
| cpeb1b | cytoplasmic polyadenylation element binding protein 1b |
| cs | citrate synthase |
| ddx46 | DEAD (Asp-Glu-Ala-Asp) box polypeptide 46 |
| dnajc22 | DnaJ (Hsp40) homolog, subfamily C, member 22 |
| dop1b | DOP1 leucine zipper like protein B |
| e2f1 | E2F transcription factor 1 |
| endou2 | endonuclease, polyU-specific 2 |
| fbxo40.1 | F-box protein 40, tandem duplicate 1 |
| fsd1 | fibronectin type III and SPRY domain containing 1 |
| gclc | glutamate-cysteine ligase, catalytic subunit |
| gins2 | GINS complex subunit 2 |
| gk5 | glycerol kinase 5 |
| gkap1 | G kinase anchoring protein 1 |
| glmna | glomulin, FKBP associated protein a |
| hddc2 | HD domain containing 2 |
| hectd3 | HECT domain containing 3 |
| hgs | hepatocyte growth factor-regulated tyrosine kinase substrate |
| hltf | helicase-like transcription factor |
| hmg20a* | high mobility group 20A |
| itgb4 | integrin, beta 4 |
| kif4 | kinesin family member 4 |
| letm2 | leucine zipper-EF-hand containing transmembrane protein 2 |
| lonp2 | lon peptidase 2, peroxisomal |
| lrrn3b | leucine rich repeat neuronal 3b |
| mapk15 | mitogen-activated protein kinase 15 |
| mlx | MAX dimerization protein MLX |
| mrpl41 | mitochondrial ribosomal protein L41 |
| nr2c2* | nuclear receptor subfamily 2, group C, member 2 |
| paip1 | poly(A) binding protein interacting protein 1 |
| pik3c3 | phosphatidylinositol 3-kinase, catalytic subunit type 3 |
| pik3r5 | phosphoinositide-3-kinase, regulatory subunit 5 |
| ppan* | peter pan homolog |
| ppp1r12a | protein phosphatase 1, regulatory subunit 12A |
| psmc5 | proteasome 26S subunit, ATPase 5 |
| qars1 | glutaminyl-tRNA synthetase 1 |
| rab24 | RAB24, member RAS oncogene family |
| rad9b | RAD9 checkpoint clamp component B |
| rfng | RFNG O-fucosylpeptide 3-beta-N-acetylglucosaminyltransferase |
| rfxank | regulatory factor X-associated ankyrin-containing protein |

| | |
|---|---|
| rgs14a | regulator of G protein signaling 14a |
| rmdn1 | regulator of microtubule dynamics 1 |
| rpl27* | ribosomal protein L27 |
| rplp0 | ribosomal protein, large, P0 |
| snrnp40 | small nuclear ribonucleoprotein 40 (U5) |
| srebf2 | sterol regulatory element binding transcription factor 2 |
| supt7l | SPT7 like, STAGA complex gamma subunit |
| taf5 | TAF5 RNA polymerase II, TATA box binding protein (TBP)-associated factor |
| tbcd | tubulin folding cofactor D |
| thoc5 | THO complex 5 |
| thtpa | thiamine triphosphatase |
| trip11 | thyroid hormone receptor interactor 11 |
| unc45b | unc-45 myosin chaperone B |
| wdr13 | WD repeat domain 13 |
| zfand4 | zinc finger, AN1-type domain 4 |
| zgc:152830 | zgc:152830 |
| zgc:162698 | zgc:162698 |
| zgc:85936* | zgc:85936 |
| zgc:92518 | zgc:92518 |

Note: Asterisks (*) indicate genes that are **not** also detected as selected in Latridae

Supplementary Table 6. Genes positively selected in Latridae [Continues next pages]

| Gene | Protein name |
|---|---|
| abhd11 | abhydrolase domain containing 11 |
| abtb1 | ankyrin repeat and BTB (POZ) domain containing 1 |
| acer1 | alkaline ceramidase 1 |
| acer3 | alkaline ceramidase 3 |
| acmsd | aminocarboxymuconate semialdehyde decarboxylase |
| aff4 | AF4/FMR2 family, member 4 |
| alkbh7 | alkB homolog 7 |
| ankzf1 | ankyrin repeat and zinc finger peptidyl tRNA hydrolase 1 |
| apex1 | APEX nuclease (multifunctional DNA repair enzyme) 1 |
| arhgap24 | Rho GTPase activating protein 24 |
| arhgap32b | Rho GTPase activating protein 32b |
| arl13a | ADP-ribosylation factor-like 13A |
| atp5f1c | ATP synthase F1 subunit gamma |
| atp6v1d | ATPase H+ transporting V1 subunit D |
| aup1 | AUP1 lipid droplet regulating VLDL assembly factor |
| babam1 | BRISC and BRCA1 A complex member 1 |
| bspry | B-box and SPRY domain containing |
| c1qbp | complement component 1, q subcomponent binding protein |
| C3H17orf75 | zgc:153240 |
| cabin1 | calcineurin binding protein 1 |
| CACFD1 | si:ch73-209e20.5 |
| capn10 | calpain 10 |
| cars2 | cysteinyl-tRNA synthetase 2, mitochondrial |
| ccdc84 | coiled-coil domain containing 84 |
| cdadc1 | cytidine and dCMP deaminase domain containing 1 |
| cdk10 | cyclin-dependent kinase 10 |
| cdkn2aip | CDKN2A interacting protein |
| cfap58 | cilia and flagella associated protein 58 |
| cmtr1 | cap methyltransferase 1 |
| cmtr2 | cap methyltransferase 2 |
| commd5 | COMM domain containing 5 |
| cpeb1b | cytoplasmic polyadenylation element binding protein 1b |
| cs | citrate synthase |
| ddx46 | DEAD (Asp-Glu-Ala-Asp) box polypeptide 46 |
| dhh | desert hedgehog signaling molecule |
| dhx29 | DEAH (Asp-Glu-Ala-His) box polypeptide 29 |
| dhx33 | DEAH (Asp-Glu-Ala-His) box polypeptide 33 |
| dhx58 | DEXH (Asp-Glu-X-His) box polypeptide 58 |
| dnajc22 | DnaJ (Hsp40) homolog, subfamily C, member 22 |
| dop1b | DOP1 leucine zipper like protein B |
| dot1l | DOT1-like histone H3K79 methyltransferase |
| e2f1 | E2F transcription factor 1 |
| e2f3 | E2F transcription factor 3 |
| ecsit | ECSIT signaling integrator |
| eif3s6ip | eukaryotic translation initiation factor 3, subunit 6 interacting protein |

| | |
|---|---|
| endou2 | endonuclease, polyU-specific 2 |
| FADS6 | fatty acid desaturase 6 |
| fam199x | family with sequence similarity 199, X-linked |
| fbxo40.1 | F-box protein 40, tandem duplicate 1 |
| fdft1 | farnesyl-diphosphate farnesyltransferase 1 |
| fgb | fibrinogen beta chain |
| flrt3 | fibronectin leucine rich transmembrane 3 |
| foxred2 | FAD-dependent oxidoreductase domain containing 2 |
| fra10ac1 | FRA10A associated CGG repeat 1 |
| fsd1 | fibronectin type III and SPRY domain containing 1 |
| gclc | glutamate-cysteine ligase, catalytic subunit |
| gins2 | GINS complex subunit 2 |
| gk5 | glycerol kinase 5 |
| gkap1 | G kinase anchoring protein 1 |
| glmna | glomulin, FKBP associated protein a |
| gpam | glycerol-3-phosphate acyltransferase, mitochondrial |
| gpat2 | glycerol-3-phosphate acyltransferase 2, mitochondrial |
| gpr135 | G protein-coupled receptor 135 |
| gpr143 | G protein-coupled receptor 143 |
| gramd1c | GRAM domain containing 1c |
| gtpbp4 | GTP binding protein 4 |
| hbegfa | heparin-binding EGF-like growth factor a |
| hddc2 | HD domain containing 2 |
| hectd3 | HECT domain containing 3 |
| herc4 | HECT and RLD domain containing E3 ubiquitin protein ligase 4 |
| hgs | hepatocyte growth factor-regulated tyrosine kinase substrate |
| higd2a | HIG1 hypoxia inducible domain family, member 2A |
| hltf | helicase-like transcription factor |
| ift22 | intraflagellar transport 22 homolog (Chlamydomonas) |
| inppl1b | inositol polyphosphate phosphatase-like 1b |
| itgb4 | integrin, beta 4 |
| IYD | si:ch211-286f9.2 |
| kbtbd3 | kelch repeat and BTB (POZ) domain containing 3 |
| kif26ba | kinesin family member 26Ba |
| kif4 | kinesin family member 4 |
| klhl30 | kelch-like family member 30 |
| kpnb3 | karyopherin (importin) beta 3 |
| krcp | kelch repeat-containing protein |
| laptm4a | lysosomal protein transmembrane 4 alpha |
| leo1 | LEO1 homolog, Paf1/RNA polymerase II complex component |
| letm2 | leucine zipper-EF-hand containing transmembrane protein 2 |
| limk2 | LIM domain kinase 2 |
| lonp2 | lon peptidase 2, peroxisomal |
| lpcat2 | lysophosphatidylcholine acyltransferase 2 |
| lrguk | leucine-rich repeats and guanylate kinase domain containing |
| lrrn3b | leucine rich repeat neuronal 3b |
| mapk15 | mitogen-activated protein kinase 15 |
| mapk6 | mitogen-activated protein kinase 6 |

| | |
|---|---|
| mcoln2 | mucolipin 2 |
| mcu | mitochondrial calcium uniporter |
| meak7 | MTOR associated protein, eak-7 homolog |
| memo1 | mediator of cell motility 1 |
| mettl13 | methyltransferase like 13 |
| miga2 | mitoguardin 2 |
| mknk1 | MAPK interacting serine/threonine kinase 1 |
| mlx | MAX dimerization protein MLX |
| mms19 | MMS19 homolog, cytosolic iron-sulfur assembly component |
| mocs3 | molybdenum cofactor synthesis 3 |
| mon1a | MON1 secretory trafficking family member A |
| mpnd | MPN domain containing |
| mrpl41 | mitochondrial ribosomal protein L41 |
| mthfr | methylenetetrahydrofolate reductase (NAD(P)H) |
| naa30 | N(alpha)-acetyltransferase 30, NatC catalytic subunit |
| ncbp2 | nuclear cap binding protein subunit 2 |
| ndufv2 | NADH:ubiquinone oxidoreductase core subunit V2 |
| nelfcd | negative elongation factor complex member C/D |
| nfat5b | nuclear factor of activated T cells 5b |
| nfrkb | nuclear factor related to kappaB binding protein |
| notch2 | notch receptor 2 |
| nploc4 | NPL4 homolog, ubiquitin recognition factor |
| nsrp1 | nuclear speckle splicing regulatory protein 1 |
| numa1 | nuclear mitotic apparatus protein 1 |
| ogfod1 | 2-oxoglutarate and iron-dependent oxygenase domain containing 1 |
| opa1 | OPA1 mitochondrial dynamin like GTPase |
| paip1 | poly(A) binding protein interacting protein 1 |
| pax8 | paired box 8 |
| pde12 | phosphodiesterase 12 |
| pdia6 | protein disulfide isomerase family A, member 6 |
| phgdh | phosphoglycerate dehydrogenase |
| phyhd1 | phytanoyl-CoA dioxygenase domain containing 1 |
| pik3c3 | phosphatidylinositol 3-kinase, catalytic subunit type 3 |
| pik3r5 | phosphoinositide-3-kinase, regulatory subunit 5 |
| plekha8 | pleckstrin homology domain containing, family A (phosphoinositide binding specific) member 8 |
| PLEKHH3 | si:ch211-18i17.2 |
| polr3e | polymerase (RNA) III (DNA directed) polypeptide E |
| pomgnt2 | protein O-linked mannose N-acetylglucosaminyltransferase 2 (beta 1,4-) |
| ppp1r12a | protein phosphatase 1, regulatory subunit 12A |
| psma1 | proteasome 20S subunit alpha 1 |
| psmc5 | proteasome 26S subunit, ATPase 5 |
| psmd8 | proteasome 26S subunit, non-ATPase 8 |
| qars1 | glutaminyl-tRNA synthetase 1 |
| qser1 | glutamine and serine rich 1 |
| rab24 | RAB24, member RAS oncogene family |
| rad9b | RAD9 checkpoint clamp component B |
| ranbp3b | RAN binding protein 3b |
| rapgef3 | Rap guanine nucleotide exchange factor (GEF) 3 |

| | |
|---|---|
| rbck1 | RanBP-type and C3HC4-type zinc finger containing 1 |
| rbm17 | RNA binding motif protein 17 |
| rfng | RFNG O-fucosylpeptide 3-beta-N-acetylglucosaminyltransferase |
| rfxank | regulatory factor X-associated ankyrin-containing protein |
| rgs14a | regulator of G protein signaling 14a |
| rhag | Rh associated glycoprotein |
| rmdn1 | regulator of microtubule dynamics 1 |
| rnf113a | ring finger protein 113A |
| rpl4 | ribosomal protein L4 |
| rplp0 | ribosomal protein, large, P0 |
| rps2 | ribosomal protein S2 |
| rps6 | ribosomal protein S6 |
| rps6kl1 | ribosomal protein S6 kinase-like 1 |
| sash3 | SAM and SH3 domain containing 3 |
| scly | selenocysteine lyase |
| sik3 | SIK family kinase 3 |
| snai1a | snail family zinc finger 1a |
| snrnp40 | small nuclear ribonucleoprotein 40 (U5) |
| snrpb | small nuclear ribonucleoprotein polypeptides B and B1 |
| snw1 | SNW domain containing 1 |
| snx24 | sorting nexin 24 |
| snx5 | sorting nexin 5 |
| srebf2 | sterol regulatory element binding transcription factor 2 |
| ssb | small RNA binding exonuclease protection factor La |
| stx18 | syntaxin 18 |
| stxbp3 | syntaxin binding protein 3 |
| suclg1 | succinate-CoA ligase, alpha subunit |
| sumf2 | sulfatase modifying factor 2 |
| supt20 | SPT20 homolog, SAGA complex component |
| supt4h1 | SPT4 homolog, DSIF elongation factor subunit |
| supt7l | SPT7 like, STAGA complex gamma subunit |
| svild | supervillin d |
| syf2 | SYF2 pre-mRNA-splicing factor |
| taf5 | TAF5 RNA polymerase II, TATA box binding protein (TBP)-associated factor |
| tbcd | tubulin folding cofactor D |
| tbrg1 | transforming growth factor beta regulator 1 |
| tex261 | testis expressed 261 |
| tfip11 | tuftelin interacting protein 11 |
| thoc5 | THO complex 5 |
| thtpa | thiamine triphosphatase |
| tm9sf4 | transmembrane 9 superfamily protein member 4 |
| tmem135 | transmembrane protein 135 |
| tmem161b | transmembrane protein 161B |
| traf7 | TNF receptor-associated factor 7 |
| trip11 | thyroid hormone receptor interactor 11 |
| ubap2l | ubiquitin associated protein 2-like |
| ufd1l | ubiquitin recognition factor in ER associated degradation 1 |
| unc45b | unc-45 myosin chaperone B |

| | |
|---|---|
| usp39 | ubiquitin specific peptidase 39 |
| usp4 | ubiquitin specific peptidase 4 (proto-oncogene) |
| usp49 | ubiquitin specific peptidase 49 |
| wasf2 | WASP family member 2 |
| wdr13 | WD repeat domain 13 |
| wdr61 | WD repeat domain 61 |
| wnt4 | wingless-type MMTV integration site family, member 4 |
| xpc | xeroderma pigmentosum, complementation group C |
| xrcc5 | X-ray repair complementing defective repair in Chinese hamster cells 5 |
| xrn2 | 5'-3' exoribonuclease 2 |
| zfand4 | zinc finger, AN1-type domain 4 |
| zgc:101663 | zgc:101663 |
| zgc:103625 | zgc:103625 |
| zgc:112294 | zgc:112294 |
| zgc:114119 | zgc:114119 |
| zgc:152830 | zgc:152830 |
| zgc:162698 | zgc:162698 |
| zgc:162879 | zgc:162879 |
| zgc:92518 | zgc:92518 |
| zwilch | zwilch kinetochore protein |

### Supplementary Figures

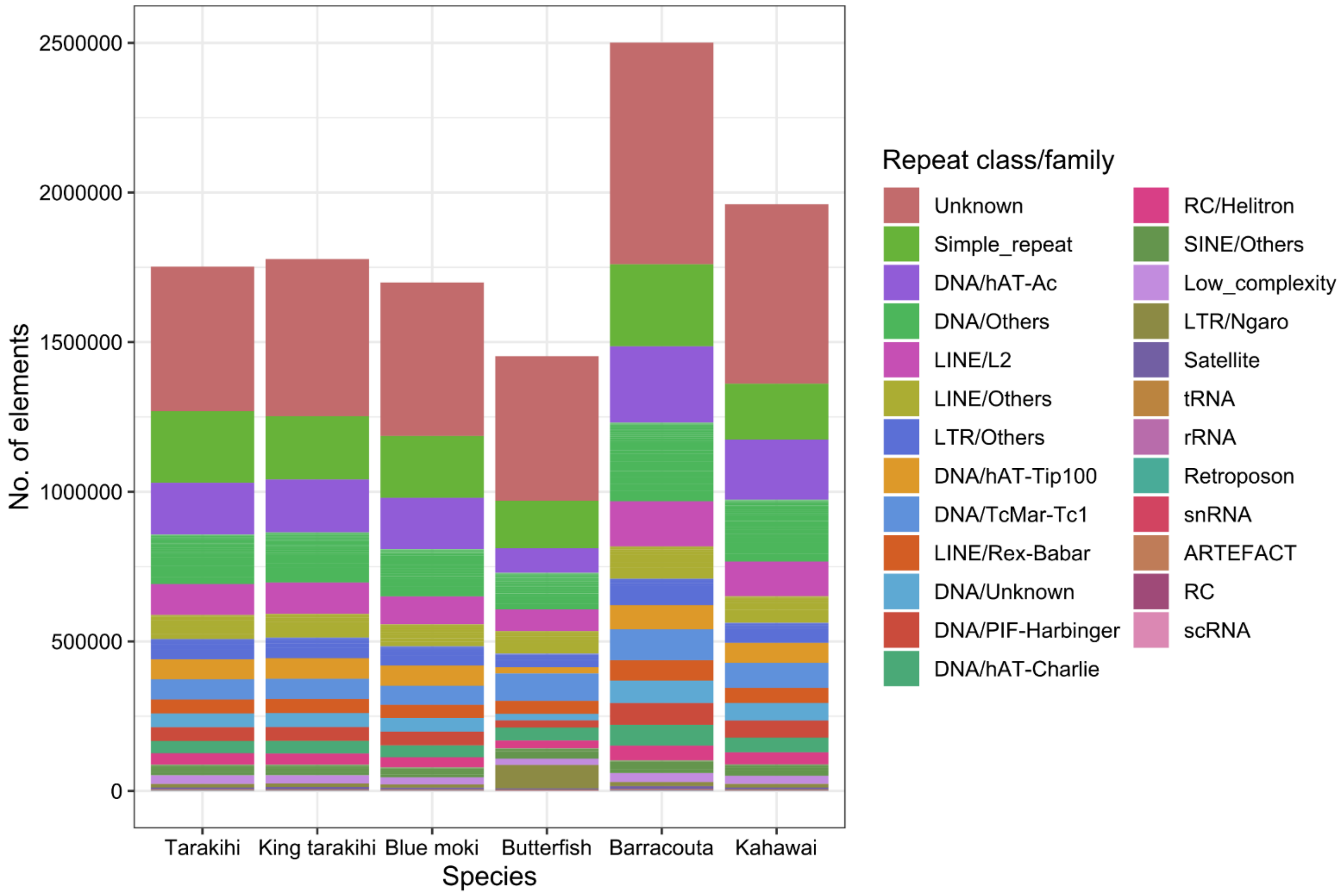


Supplementary Figure 1. Proportions of the most represented families of repeat elements in the genomes. “Other” includes all the families that are not in the top ten of the most abundant RE in at least one species. Repeat elements families are sorted vertically based on their abundance in the tarakihi.

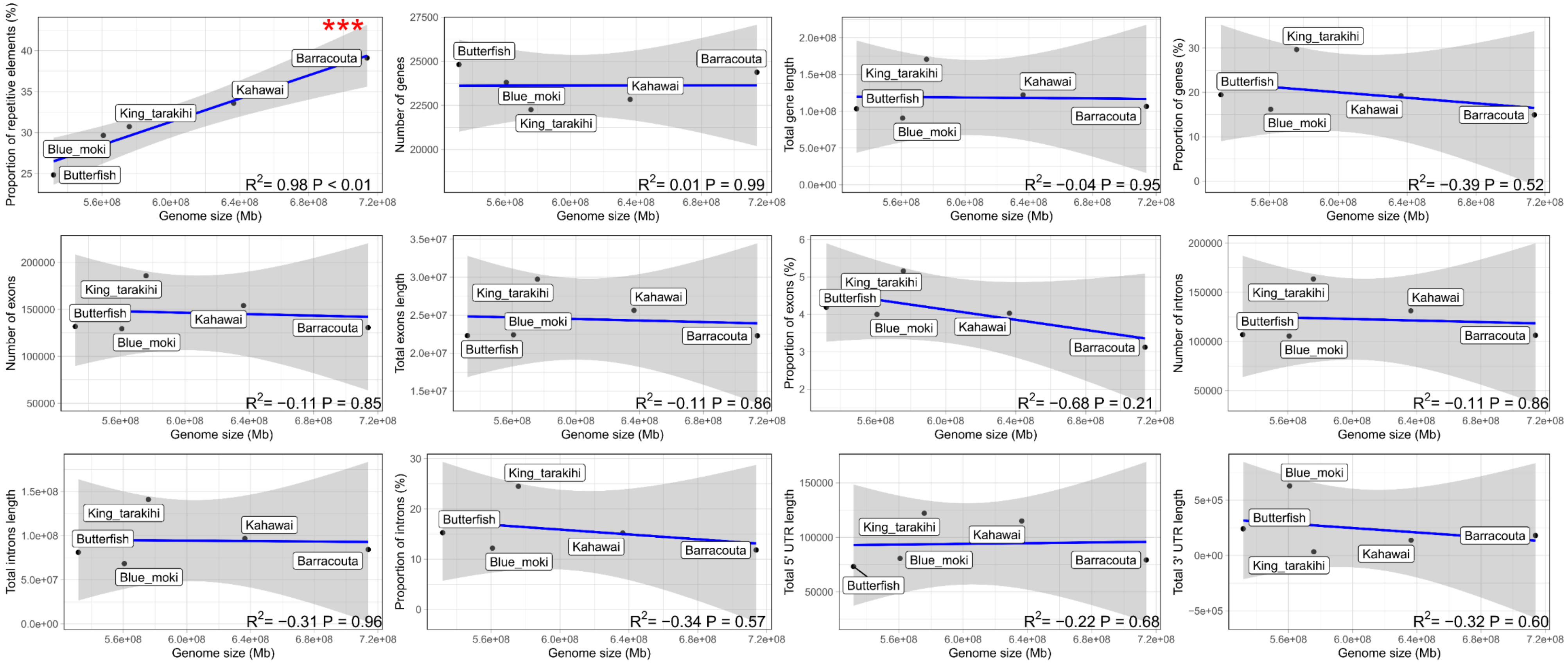


Supplementary Figure 2. Correlation between genome size and proportion, number and length of genomic features annotated in the genome assemblies without including tarakihi, with corresponding Pearson correlation coefficient (R2) and p-value (P). Grey area is the 95% confidence interval. Red asterisks indicate significance, with P ≤ 0.1 (*), 0.05 (**), and 0.01 (***).
